# Dynamics of *bicoid* mRNA localisation and translation dictate morphogen gradient formation

**DOI:** 10.1101/2024.11.11.622966

**Authors:** T. Athilingam, E.L. Wilby, P. Bensidoun, A. Trullo, M. Verbrugghe, X. Shi, M. Lagha, T.E. Saunders, T.T. Weil

## Abstract

The transcription factor Bicoid (Bcd) protein guides early *Drosophila* patterning and is the best-characterised morphogen. The source of the morphogen, *bcd* mRNA, is maternally deposited during oogenesis and localised to the anterior pole of the mature oocyte. While the spatiotemporal interpretation of the Bcd morphogen gradient has been intensely studied, when and where Bcd protein is produced and how this protein gradient is dynamically shaped remains contentious. Here, we use the SunTag reporter system to quantitatively examine the spatiotemporal profile of *bcd* mRNA translation *in vivo*. We show that association with Processing bodies (P bodies) in mature oocytes prevent premature *bcd* mRNA translation. Following egg activation, *bcd* mRNA dissociates from P bodies and translation is observed at the anterior pole. Translation remains restricted to the anterior domain throughout early development, even after nuclear migration in the syncytial blastoderm. At cellularisation, translation ceases and the remaining *bcd* mRNA associates with reformed P bodies, which appear to block any further translation. We use these observations to create a new modified source-diffusion-degradation model of Bcd gradient formation that has spatiotemporally varying production. Overall, our study reveals that *bcd* mRNA translation is tightly controlled in space and time during oogenesis and early embryogenesis.

## Introduction

The concept of signalling molecules, coined morphogens, was proposed by Turing (1). These molecules help guide spatial patterning by varying in concentration across developing systems. Downstream target genes respond in a concentration-dependent manner to the morphogen gradient (2). Bicoid (Bcd) protein is a transcription factor found in the early *Drosophila* blastocyst and was the first morphogen gradient identified (3). A concentration gradient of Bcd extends along the anterior-posterior (AP) axis from highest at the anterior pole of the embryo and reaches around 70% of the embryo’s length. The expression of target genes, like *hunchback,* depends on the local Bcd concentration (4–6).

Shortly after its discovery, the Synthesis, Diffusion, Degradation (SDD) model was introduced to explain how the Bcd gradient forms (3, 7). This model assumed that Bcd is produced at the anterior pole, then diffuses through the embryo and is subsequently degraded. The model predicted a concentration profile that decreased exponentially in steady state, consistent with *in vivo* observations (7–9). There are several mechanisms that can create morphogen profiles that decay exponentially (10, 11). Alternatives to the SDD model include: (i) local production of Bcd protein by *bcd* mRNA spatially distributed along the embryo (12); (ii) early translation of *bcd* in the oocyte which would allow the Bcd morphogen to spread quickly before patterning happens in later nuclear cycles (13); and (iii) nuclear dilution, which does not require protein degradation (14). These models predict different times and places for *bcd* mRNA translation. Still, they can all create exponential-like concentration gradients under the right conditions and time scales for patterning (15–17).

Localised *bcd* mRNA first appears at the anterior region in mid-stage oocytes (18–20). As oogenesis progresses, *bcd* transitions from a continual active transport to stable anchoring at the anterior cortex (20, 21). By the end of oogenesis, the mature oocyte (stage 14) is endowed with all the *bcd* needed for anterior patterning in the embryo (18–21). There is no biochemical evidence that *bcd* translation happens before egg activation (22–24), a conserved process that triggers several major changes in the egg (25–28). These changes include restarting meiosis, altering the translational landscape, and reconfiguring the cytoskeleton. These events occur as the mature egg moves into the oviduct before fertilisation in the uterus (29). Translational repression of *bcd* in the oocyte is proposed to occur through its association in a ribonuclear protein (RNP) granule, a class of biomolecular condensates (30, 31).

Most research on biomolecular condensate composition, physical properties, and function has focused on single cell models and *in vitro* systems (32–36). This provides an excellent basis for studying ribonucleoprotein (RNP) granule physiology and translational regulation in multicellular organisms. Genetic and physical manipulation in *Drosophila* have recently helped us understand the biological relevance of RNP granules in a developmental context (37–41). For example, visualising *nanos* mRNA translation shows that germ granules can either be sites of translation or repression depending on the location of the mRNA (40, 41). Also, the germ-line determinant *oskar* (*osk*) mRNA associates with “solid-like” condensates where its unique secondary structure is required to maintain *osk* mRNA inside the condensates, where it is translationally repressed (37).

A similar model of mRNA regulation via association with an RNP granule had previously been proposed for *bcd* mRNA (30, 31). Once localised to the anterior, *bcd* is sequestered in the core of Processing bodies (P bodies) away from ribosomes (30). P bodies are an evolutionarily cytoplasmic RNP granule. Depending on the cellular context, they are implicated in storing and degrading mRNA (32, 42). A variety of proteins that help regulate mRNA are enriched in P bodies including the DEAD-box family RNA helicase, Maternal Effect at 31B (Me31B). It has homologies with proteins in other species: DDX6 in humans, Dhh1 in *S.cerevisiae*, Xp54 in *Xenopus*, and Cgh-1 in *C.elegans* (30, 43). Previous work has shown that Me31B functions both in repression and clearance of maternal transcripts in early *Drosophila* embryogenesis (44).

Previous studies on the Bcd morphogen gradient have mainly looked at two areas. One focuses on the regulation of *bcd* mRNA during oogenesis (30, 45–47). The other examines Bcd protein dynamics after translation (7, 10, 17, 48, 49). To fully understand how the Bcd gradient forms, we need to integrate these two approaches. Here, we used the SunTag/scFv imaging-based reporter system (50–57) to quantitatively assay *bcd* translation in *Drosophila* oocytes and embryos. We show that *bcd* mRNA is not translated when it is in P bodies at the end of oogenesis. Following egg activation, or experimentally driven disassociation, *bcd* is no longer in P bodies and is translated. We observed *bcd* translation starts quickly after release from P-bodies, but it happens only at the anterior pole of the embryo. After nuclear migration, *bcd* localisation appears dispersed though the location of translation remains unchanged. We saw a decrease in *bcd* translation in nuclear cycle (n.c.) 11 onwards, consistent with observations from assaying the protein levels (58). Motivated by our experimental observations, we demonstrate that a modified SDD model, with spatially and temporally varying Bcd production, is consistent with our results.

## Results

### *bcd* translation begins after egg activation

To test when and where *bcd* mRNA is translated, we generated a *SunTag-bcd* transgene, containing an array of 32 SunTag GCN4 repeats (*bcdSun32*) (Figure 1A, Methods). We first validated our reporter in early embryos, where Bcd protein is known to be present (59). We performed fluorescence single molecule *in situ* hybridisation (smFISH) to label *bcdSun32* and antibody labelling to visualise the SunTag nascent peptide (Figure 1B). We verified the *bcdSun32* reporter by showing two populations of *bcdSun32* particles: *bcdSun32* mRNA particles co-localizing with the GCN4 antibody signal, (implying polysome and mRNA engaged in translation); and *bcdSun32* mRNA molecules that do not overlap with anti-GCN4 (not in translation) (Figure 1B). We also performed this validation using the fluorescent detector, the single-chain antibody scFV, which is genetically encoded and maternally provided (*nos-scFv-MSGFP2*) (60). To further test the reporter, we expressed *bcdSun32* in oocytes. We observed no translation in oocytes (Figure 1C-D) which matches what we predicted from earlier findings of *bcd* mRNA translational regulation (61, 62).

**Figure 1:**
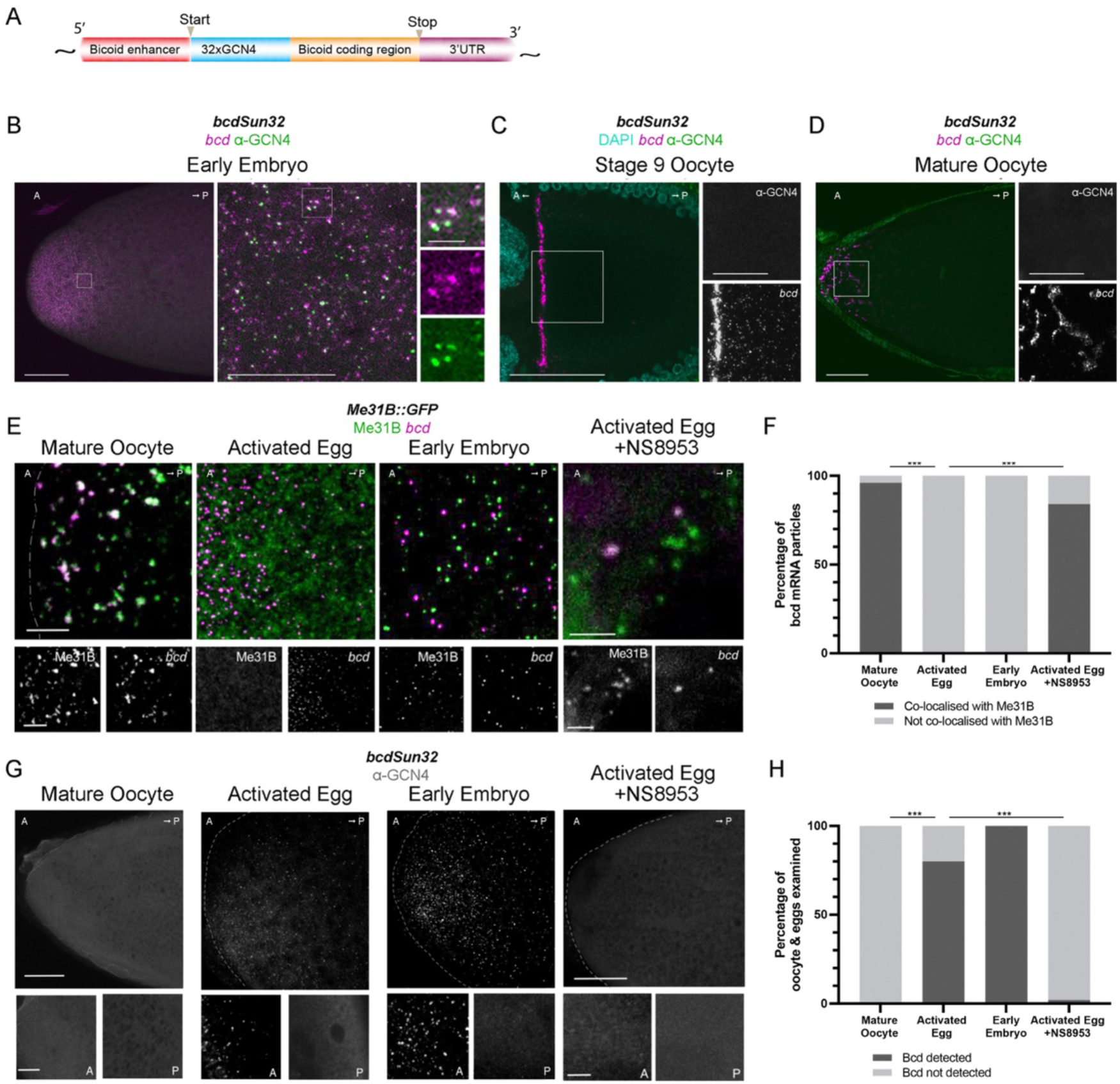
*bcdSun32* reporter enables visualisation of *bcd* mRNA translation. A: Schematic illustrating the insertion of 32x SunTag repeats at the N-terminal of the *bcd* coding region after the start codon. B: Representative image of *bcdSun32* in the early embryo (from 15 samples) with *bcd* translation detected by immunostaining for GCN4 (green) and *bcd* mRNA (magenta). Boxed region magnified on right. Boxed region on right further magnified. N = 20. Scale bars: left 50 µm; middle 10 µm; right 3 µm. C-D: Representative fixed 5-10 µm Z-stack images showing *bcdSun32* labelled *bcd* (magenta, smFISH) localising to the anterior margin of stage 9 and 14 (mature) oocytes. SunTag labelled Bcd protein (green, anti-GCN4) is not detected in stage 9 or 14 oocytes. N = 20. Scale bars: 50 µm; zoom 10 µm. E: Representative fixed 10 µm Z-stack images (from 10 samples) showing Me31B::GFP (green, anti-GFP-nanobody) and *bcd* (magenta, smFISH) at the anterior in mature oocyte, *in vitro* activated egg, early embryo 30-minute post fertilisation, and *in vitro* activated egg incubated with NS8953, a calcium channel blocker. Single channel images are shown for Me31B and *bcd*. Scale bar: 5 μm. F: Quantification of the co-localisation (as shown in E) between *bcd* particles and Me31B::GFP particles. N = 6 for mature oocytes, *in vitro* activated egg, early embryos. N = 3 for *in vitro* activated egg incubated with NS8953 due to difficulty with particle detection post oocyte swelling. G: Representative fixed 10 µm Z-stack images (from 10 samples) showing BcdSun32 protein (anti-GCN4) at the anterior of a mature oocyte, *in vitro* activated egg, early embryo 30-minute post fertilisation, and *in vitro* activated egg incubated with NS8953. Zoomed images of the mature oocyte show that Bcd protein is not detected at the anterior posterior pole of the same mature oocyte or *in vitro* activated egg incubated with NS8953. However, zoomed images of the *in vitro* activated egg and early embryo 30-minute post fertilisation show that BcdSun32 protein was detected at the anterior and not the posterior pole. Scale bar: 20 μm; zoom 2 μm. H: Quantification of the number of mature oocytes or early embryos that displayed *bcd* translation as measured by the presence of BcdSun32 protein at the anterior but not the posterior. N = 10 for each condition. Anterior to the left and posterior to the right in all images.

Previous work on P bodies has shown they do not contain ribosomes and can act as sites of mRNA storage (63–65). To test if *bcd* mRNA is translationally repressed when in P bodies, we calculated the amount of *bcd* mRNA co-localisation to P bodies. We focused on the developmental transition from the mature egg to the early embryo. Our previous qualitative data postulated that location in P bodies is key for controlling *bcd* translation (31). Using single molecule fluorescence *in situ* hybridisation (smFISH) and the standard P body marker Me31B, we confirmed that *bcd* is in P bodies before egg activation (Figure 1E-F). Then, during the transition from a mature oocyte to activated egg, P bodies disassembled and *bcd* was no longer co-localised with P bodies (Figure 1E-F) (30, 31). We also confirmed that *bcd* becomes dispersed, losing its tight association with the anterior cortex (Figure 1E) (31). In the early embryo, the distribution of *bcd* stayed the same while ‘embryonic’ P bodies formed (Figure 1E). There was no co-localisation detected between *bcd* and P bodies at this stage, consistent with past work showing that the re-formed P bodies have different traits compared to those in the oocyte (Figure 1F) (31).

We next show that disrupting the calcium wave in the activated egg caused a failure to disperse P bodies during *ex vivo* egg activation (Figure 1E). In these activated eggs, *bcd* mRNA remains co-localised with the retained P bodies (Figure 1F). While addition of the chemical NS8593 disrupts the calcium wave, other phenotypic markers of egg activation are still observed (58). Based on these results and previous observations (31, 66), we hypothesised that the loss of co-localisation between *bcd* and P bodies correlates with *bcd* translation.

To test this model, we used the *bcdSun32* to observe translation during this transition. Before egg activation, Bcd protein was not detected (Figure 1G-H). However, after egg activation, *bcd* mRNA translation occurred rapidly and continued after egg laying (25, 67) (Figure 1G-H). Consistent with previous findings, we only observed *bcd* translation at the anterior of the activated egg and early embryo (Figure 1G-H) (3, 68). Furthermore, we did not see *bcd* translation in NS8593 treated activated mature eggs (Figure 1G-H). This means that the P bodies likely serve as a barrier, keeping *bcd* transcripts within a translation-free environment in the oocyte.

### Premature release of *bcd* mRNA results in ectopic translation

Altering the physical state of an RNP granule can impact its ability to store mRNA (31, 37, 69). To further test if *bcd* storage in P bodies is required for translational repression, we monitored mRNA association with P bodies after chemical treatment known to disrupt P body integrity (31, 70–72). After treating mature oocytes with these solutions, we observed BcdSun32 protein in the oocyte anterior (Figure 2A-B). We checked the treated mature eggs for egg activation markers to ensure that the chemical treatment did not trigger activation, which could cause translation (73). Chemically treated eggs failed the activation test of bleaching for two minutes (74). This confirms that the storage in P bodies is crucial to maintain *bcd* translational repression (31).

**Figure 2:**
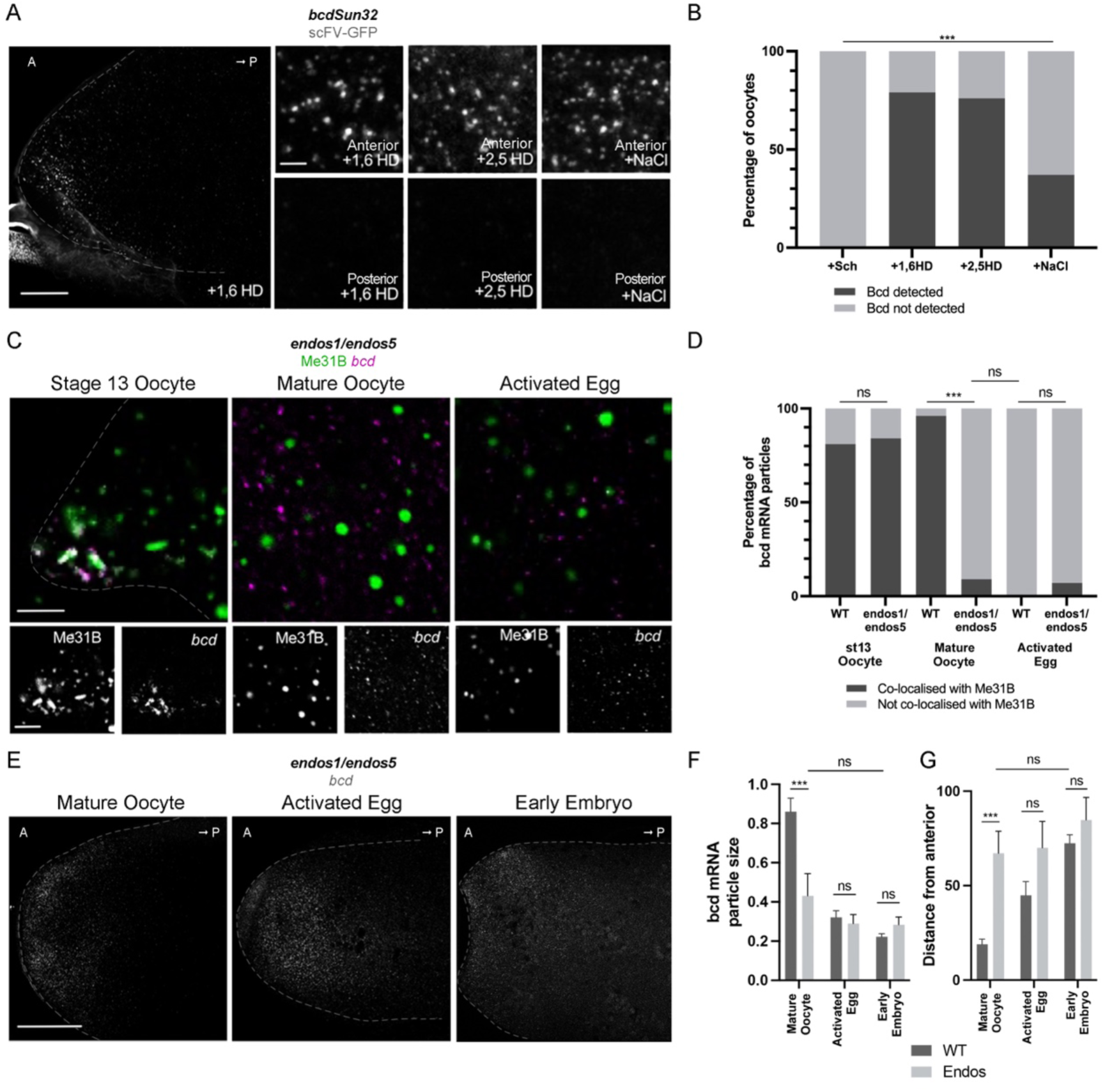
Chemical and genetic manipulation of P bodies can cause *bcd* mRNA misregulation. A: Representative fixed 10 µm Z-stack image (from 10 samples) showing the overall distribution of BcdSun32 protein (scFV-GFP, anti-GFP-nanobody) at the anterior of a stage 14 (mature) oocyte incubated in 5% 1,6 hexanediol for 20 minutes. Zoomed images show that BcdSun32 protein is detected at the anterior pole and not at the posterior pole of the same 5% 1,6 hexanediol, 5% 2,5 hexanediol, and 400 mM NaCl treated mature oocytes. Scale bars: Left 20 μm; right 2 μm. B: Quantification of experiments shown in A. The number of oocytes that displayed Bcd protein at the anterior as measured by the presence of BcdSun32 at the anterior of the oocyte, but not the posterior. N = 30 oocytes for each treatment. Scale bar: 5 μm. C: Representative fixed single plane image showing the distribution of Me31B::GFP (green, anti-GFP-nanobody) and *bcd* (magenta, smFISH) in an Endos mutant stage 13 oocyte, mature oocyte, and an *in vitro* activated egg. Scale bar: 5 μm. D: Quantification of experiments shown in C. Colocalisation is shown between *bcd* and Me31B in both wildtype and mutant stage 13 oocytes and in wildtype stage 14 oocytes. Significantly less, or no colocalisation is detected in mutant stage 14 oocytes or both wildtype and mutant activated eggs. N = between 6-10 oocytes for each genotype. E: Representative fixed 10 µm Z-stack image showing the overall distribution of *bcd* (smFISH) in an Endos mutant background at the anterior of the mature oocyte, *in vitro* activated egg, and early embryo. *bcd* appears to be particulate and diffuse at the anterior pole of all samples. Scale bars: 20 μm. F: Quantification of experiments shown in E. Quantification of the size of *bcd* particles in Endos mutant background compared to the wildtype in mature oocytes, activated eggs, and early embryos. Particles in wildtype mature oocytes are significantly larger than in Endos mutant backgrounds. Particles in an Endos mutant background all of a similar size and are not significantly different from particles in wildtype activated eggs or early embryos. Wilcoxon signed-rank test, *** *p* < 0.001, N = 5. G: Quantification of experiments shown in E. Quantification of the distance from the anterior of *bcd* particles in an Endos mutant background compared to the wildtype in mature oocytes, activated eggs, and early embryos. In mature oocytes, particles in an Endos mutant background are significantly further from the anterior than in wildtype. Particle distance from the anterior is not significantly different Endos mutant mature oocytes as compared to wildtype early embryos. Wilcoxon signed-rank test, *** *p* < 0.001, N = 5. Anterior to the left and posterior to the right in all images.

To further test this model, we used mutations in *Drosophila* Endosulfine (Endos), which is part of the conserved phosphoprotein ⍺-endosulfine (ENSA) family (75). This caused a change in RNP granule phenotype after oocyte maturation, similar to that observed with chemical treatment (Figure 2C) (31). This temporal effect matched the known roles of Endos as the master regulator of oocyte maturation (75, 76). *endos* mutant oocytes lost the co-localisation of *bcd* and P bodies, concurrent with P bodies becoming less viscous during oocyte maturation (Figure 2D, Figure S1). Particle size and position analysis showed that *bcd* mRNA prematurely exhibits an embryo distribution in these mutants (Figure 2E). Due to genetic and antibody constraints, we are unable to test for translation of *bcd* in the *endos* mutant. However, it follows that *bcd* observed in this diffuse distribution outside of P bodies would be translationally active (Figure 2E-F).

### *bcd* mRNA translation is spatially restricted during early *Drosophila* embryogenesis

We next addressed when and where the embryo translates *bcd* mRNA, if *bcd* translation is limited to the anterior pole, and when does the maximum translation occur. We considered that the size of *bcdSun*32 may impede mRNA biogenesis and the function of the protein as a transcription factor. Therefore, we generated a second transgene with ten GCN4 repeats (*bcdSun10*) (Figure S2). Importantly, *bcdSun10* rescued the *bcd*^E1^ null mutant suggesting that the tag does not greatly affect function (Figure S2). Immuno-smFISH for GCN4 and *bcd* revealed translating and non-translating *bcd* particles in the anterior pole of the embryo (Figure 3). Individual *bcd* particles (inferred from diffraction limited spots) colocalised with varying intensities of translating bcd-SunTag nascent proteins (Figure S3). At n.c.4, we saw a strong link between *bcd* mRNA and translation sites in the anterior, with most translation events happening within 100 μm of the anterior pole on the dorsal surface, and no translation observed more than 100 μm from the anterior pole (Figure 3B-C). The Bcd protein gradient is known to have small but reproducible differences between dorsal and ventral surfaces (4). Consistent with this, the translation profile differed between the dorsal to ventral sides, likely due to the asymmetric *bcd* spread on the DV-axis and on their surfaces (Figure 3A-A’’ lower panels, Figure S4).

**Figure 3:**
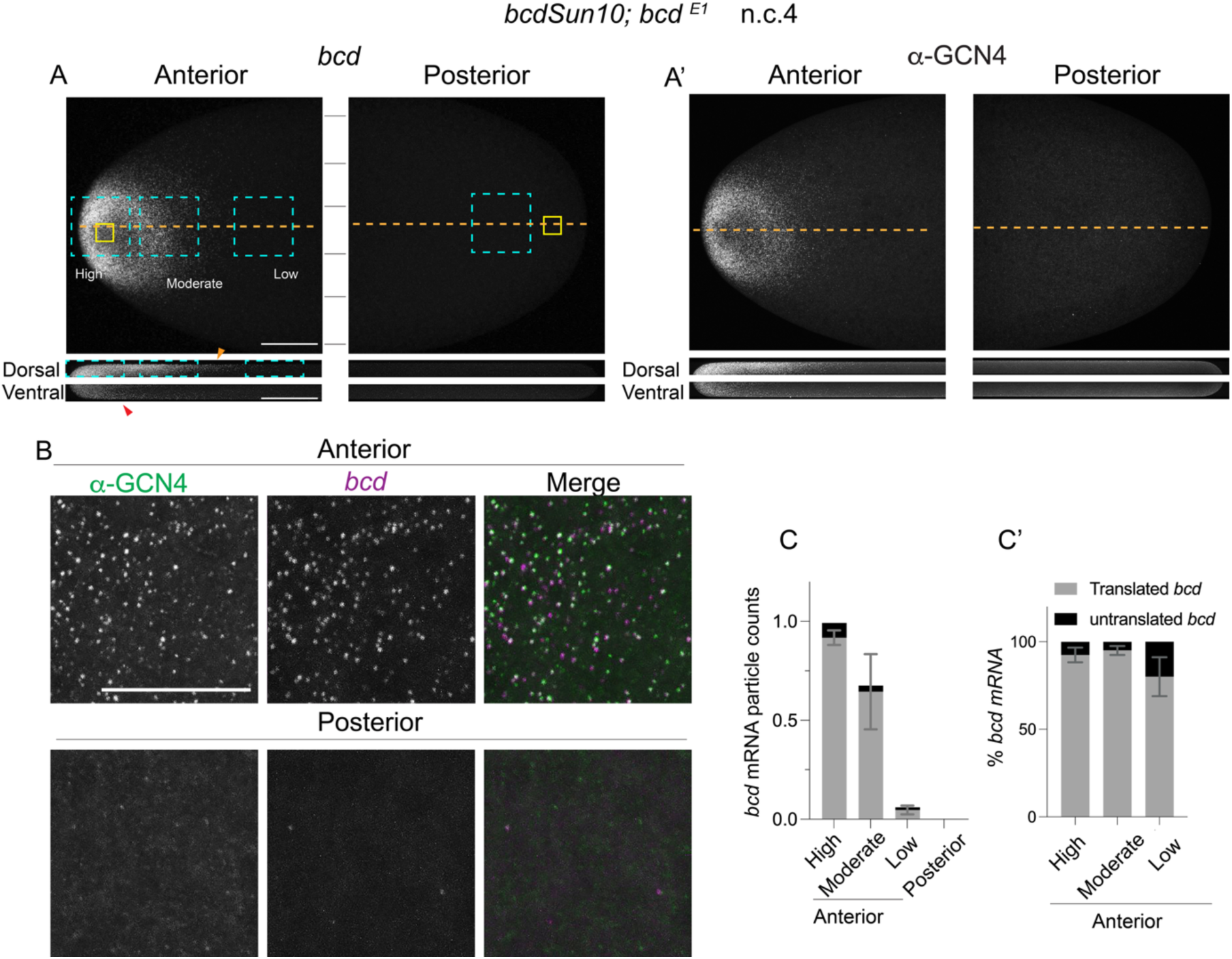
*bcd* mRNA is translated exclusively in the anterior domain of the early *Drosophila* embryo. A-Aʹ: Projected Z-stacks of the anterior and posterior domain of n.c. 4 embryo expressing the *bcdSun10* transgene, stained for *bcd* coding region (magenta) and SunTag, anti-GCN4 (green) using Immuno-FISH. The projected cross sections in the bottom panel show the dorsal and ventral surface of the anterior and posterior domains of the embryo. Translating *bcdSun10* is shown in A’ along with projected cross sections. Scale Bars: 50 µm. B: Yellow boxes from A are magnified to show colocalisation of SunTag to *bcd*. Scale Bars: 10 µm C: Normalised *bcd* counts from the cyan boxes in A located at regions of varied mRNA counts across AP-axis. Data normalised to the total mRNA counts from the most anterior region analysed. N = 3 embryos. Cʹ: Percentage of translated and untranslated *bcd* mRNA at different locations (cyan boxes in A) in the anterior and posterior domains (N = 3 embryos) in n.c. 4. Error bars = s.d.

We next quantified the number of translation events in different regions of n.c. 4 embryos (Methods, Figure S3). In the anterior region of the dorsal surface, less than 50 µm from the pole, above 90% of *bcd* particles are engaged in translation (Figure 3C-C’). Over 98% of all observed translation events occurred within 75 µm of the anterior pole on the dorsal side, with almost none seen beyond 100 µm as the *bcd* numbers are negligible. Overall, we see consistent translation efficacy irrespective of spatial differences. We conclude that during early embryogenesis, before nuclear migration to the cortex, *bcd* translation only occurs in the anterior region and mirrors the anterior-localised *bcd* distribution.

### Spatiotemporal pattern of *bcd* mRNA translation

How does translation of *bcd* vary during embryogenesis? We analysed 3D image stacks at different stages of embryo development from n.c. 4 through n.c. 14. Before n.c.9, there was sustained translation of *bcd* in a region near the anterior cortex (Figure 4A). After nuclear migration in n.c. 9, we observed a gradual decrease in translation events and by the end of n.c. 14, translation events were scarcely detected (Figure 4A-B). To test if the efficiency of translation varied at different positions in the embryo, we quantified and compared the intensity of translation events of single *bcd* particles in the anterior and posterior domain. We did not see any clear changes in the translation intensity between these locations (Figure 4C).

**Figure 4:**
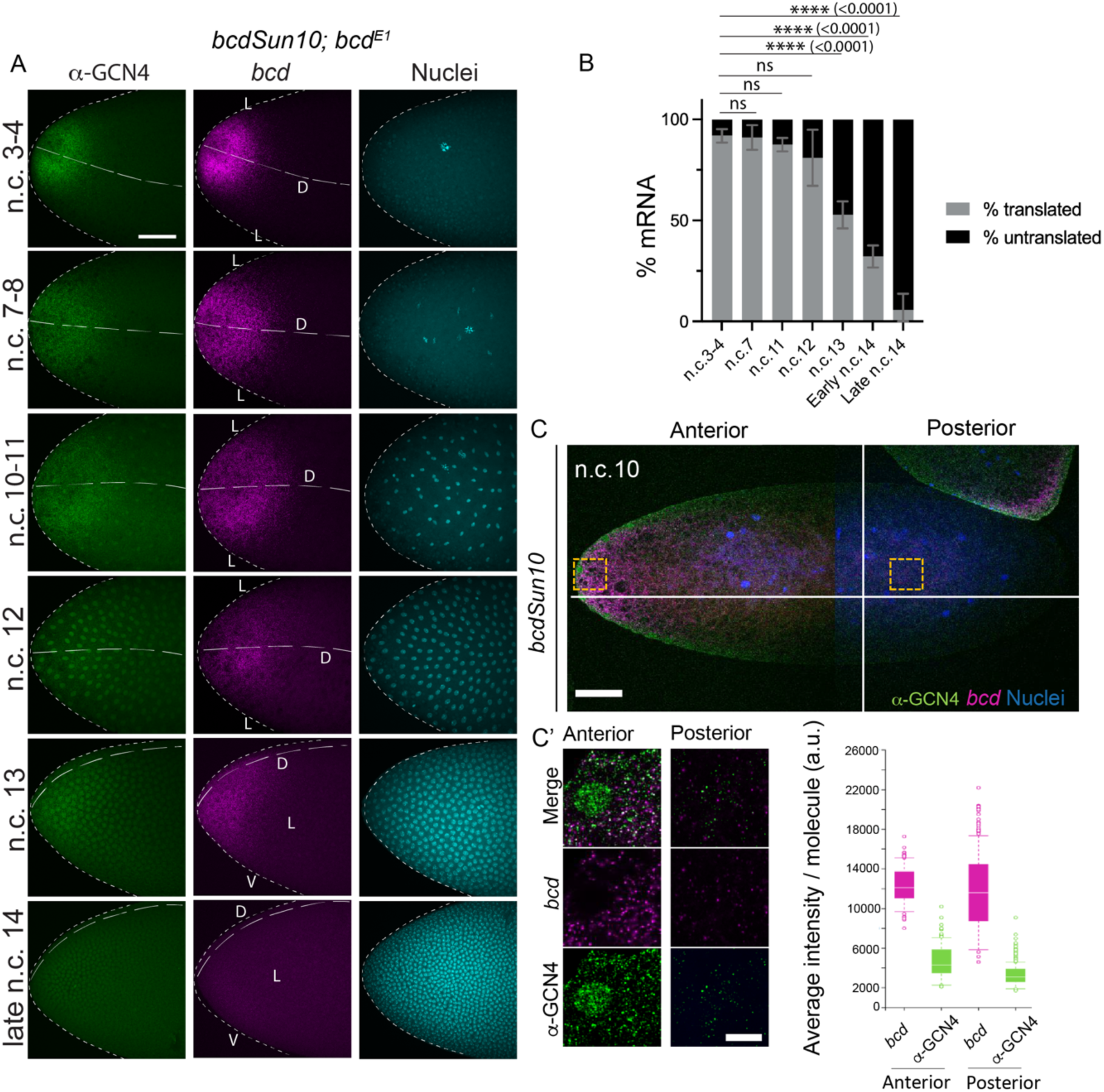
Temporal changes in *bcd* translation from early to late blastoderm. A: *bcdSun10* translation detected through immuno-FISH marked for SunTag translation (anti-GCN4, green) and *bcd* (magenta) from the early to late blastoderm embryo. D = dorsal; V = ventral; V = ventral. Scale bars: 50µm. B: Percentage of *bcd* undergoing translation at different nuclear cycles. N = 3 embryos in each of the nuclear stages. Error bars = s.d. C: Projected Z stacks of the anterior and posterior domains embryos expressing *bcdSun10* transgenes. *bcd* reporters and translation spots were detected by Immuno-FISH using *bcd* probes and anti-GCN4 shown in magenta and green respectively. Quantification of the single molecule average intensity for *bcd*-reporters smFISH and GCN4 immunofluorescence signals in anterior and posterior domains (lower right panel), N=3 embryos. Scale bars: C 50 µm; Cʹ 10 µm.

Translation can depend on the location of mRNA within the cytoplasm. In the early *Drosophila* embryo, for example, apical *twist* mRNA has lower translation than basal mRNAs (55). We analysed and compared the translating *bcd* particles above and below the cortical nuclei from n.c. 9 to late n.c.14. We did not see any significant differences in the translation of *bcd* based on their position, however, there appears an enhanced translation of *bcd* localised basally to the nuclei (Figure S5). We also noticed that upon migration, *bcd* localisation at the anterior pole became more dispersed (Figure S6A). The interphase nuclei and their subsequent mitotic divisions appeared to displace *bcd* towards the apical surface (Figure S6B).

Our *bcdSun10* not only marks the translating nascent Bcd-SunTag but can also generate a long-range, functional gradient. We can directly test different models of Bcd protein production (3, 10, 12, 58) and how this impacts the formation of the Bcd morphogen gradient. We first compared the concentration profiles of *bcd* mRNA and the protein distribution in our *bcdSun10* line from n.c. 1-8 (prior to cortical nuclear migration), n.c. 9-13 (post cortical nuclear migration), and during n.c.14 (leading up to cellularisation) (Figure 5A). We noticed an overall spread of BcdSun10 beyond the *bcd* signal, consistent with an SDD-like model for gradient formation (Figure 5B-D). The extent of *bcd* mRNA was localised to the anterior in the early embryo, with some dispersal post cortical nuclear migration (Figure 5C). However, there was no apparent long-ranged *bcd* mRNA gradient. For BcdSun10, the early protein gradient was steeper than later in the blastoderm, and always with further extent than the *bcd* mRNA distribution (Figure 5D). We saw a reduction of Bcd intensity in n.c. 14, consistent with previous reports (Figure 5E) (77).

**Figure 5:**
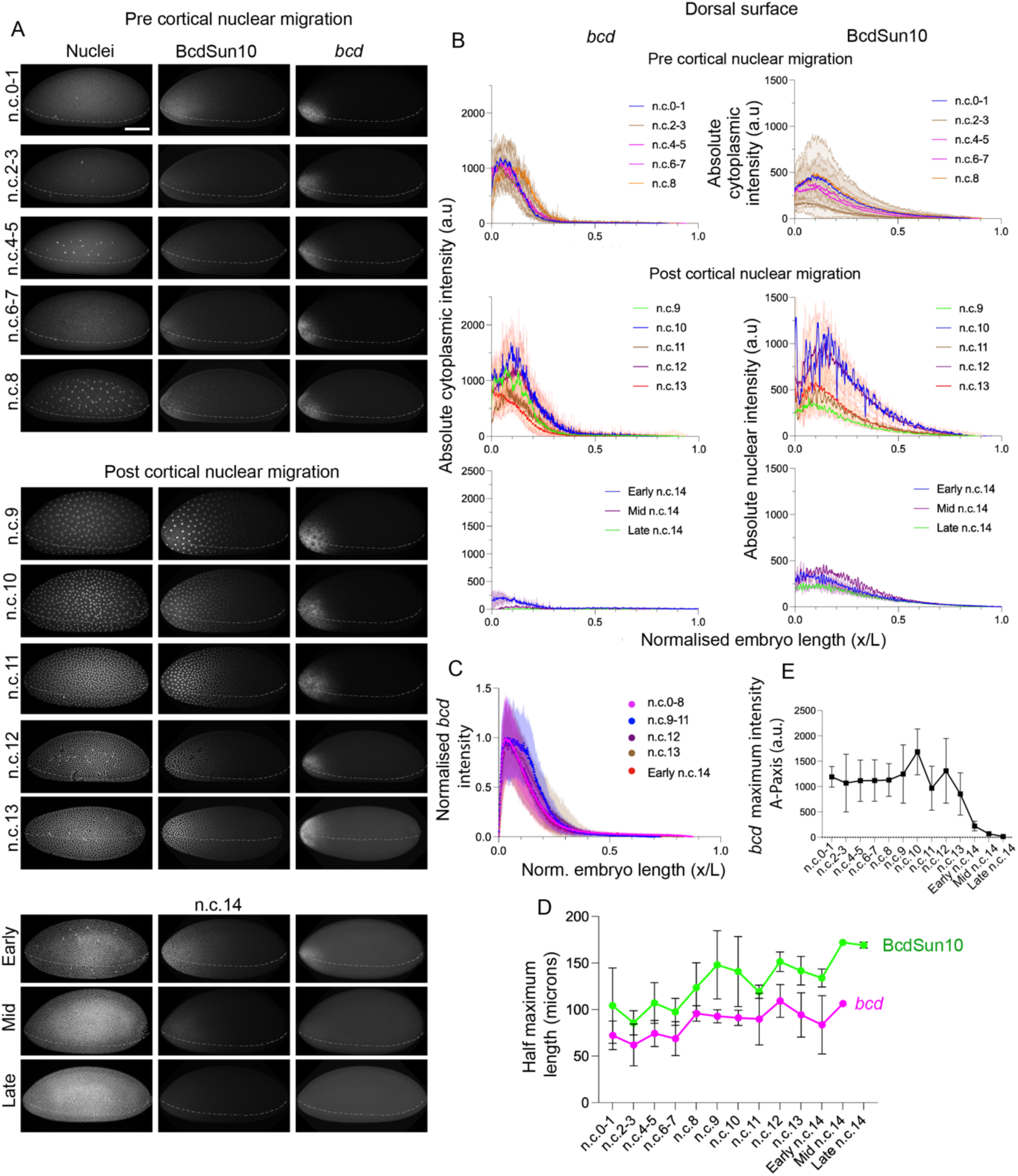
The *bcdSun10* gradient is formed by spread from the translation domain. A: *bcdSun10;bcd^E1^*embryos at different stages, showing the evolution of the SunTag-Bcd gradient (middle panel) in pre-cortical, post-cortical nuclear migration and n.c.14 with respect to the *bcd* distribution (right panel). The embryos are stained with anti-GCN4 antibody (Bcd-SunTag), *bcd* (mRNA) and DAPI (nuclei). Embryos classified based on the nuclear counts and internuclear distances from the DAPI staining. Broken lines indicate the mid-dorsal lines of the embryo. Scale bar: 50 µm. B: Line plots showing the intensity of *bcd* (left) and anti-GCN4 (right) measured across the AP-axis of the embryo’s mid dorsal surface at different blastoderm stages. C: Line plot showing the intensity of *bcd* across the AP-axis of the embryo’s mid dorsal surface. D: Distance from anterior pole for *bcd* maximum intensity to decrease by half from peak value in anterior. E: Maximum intensity *bcd* across cycles. C-E, embryo numbers: N = 3 (n.c.0-1), N = 4 (n.c.2-3), N = 5 (n.c.4-5), N = 3 (n.c.6-7), N = 3 (n.c.8), N = 3 (n.c.9), N = 3 (n.c.10), N = 2 (n.c.11), N = 4 (n.c.12), N = 6 (n.c.13), N = 2 (Early n.c.14), N = 1 (Mid n.c.14), and N = 2 (Late n.c.14).

Overall, we observe a steady shift in the domain of *bcd* mRNA translation, which correlates with nuclei movements. However, this shift does not result in substantially more dispersed *bcd* translation along the AP-axis. More than 98% of all translation events occurred within 75μm of the anterior pole, regardless of the nuclear cycle. These results support a model of localised Bcd production, restricted to a domain near the anterior pole.

### *bcd* mRNA degradation in n.c. 14 embryos

We showed a significant reduction in *bcd* mRNA translation from n.c. 13 onwards, and smFISH confirms that during n.c. 14 *bcd* is degraded (Figure 6A-C) (12, 78). Previous work has shown mRNA co-localisation with embryonic P bodies is correlated with degradation (79–81). We next tested if *bcd* was targeted for degradation in embryonic P bodies. We observed *bcd* and Me31B at early, mid, and late n.c. 14 (Figure 6Aʹ-Cʹ). With smFISH, we showed that *bcd* was evenly distribution on both the apical and basal sides of the nuclei. Co-localisation analysis revealed that *bcd* is not in P bodies at early n.c. 14 (Figure 6Aʹ, D).

**Figure 6:**
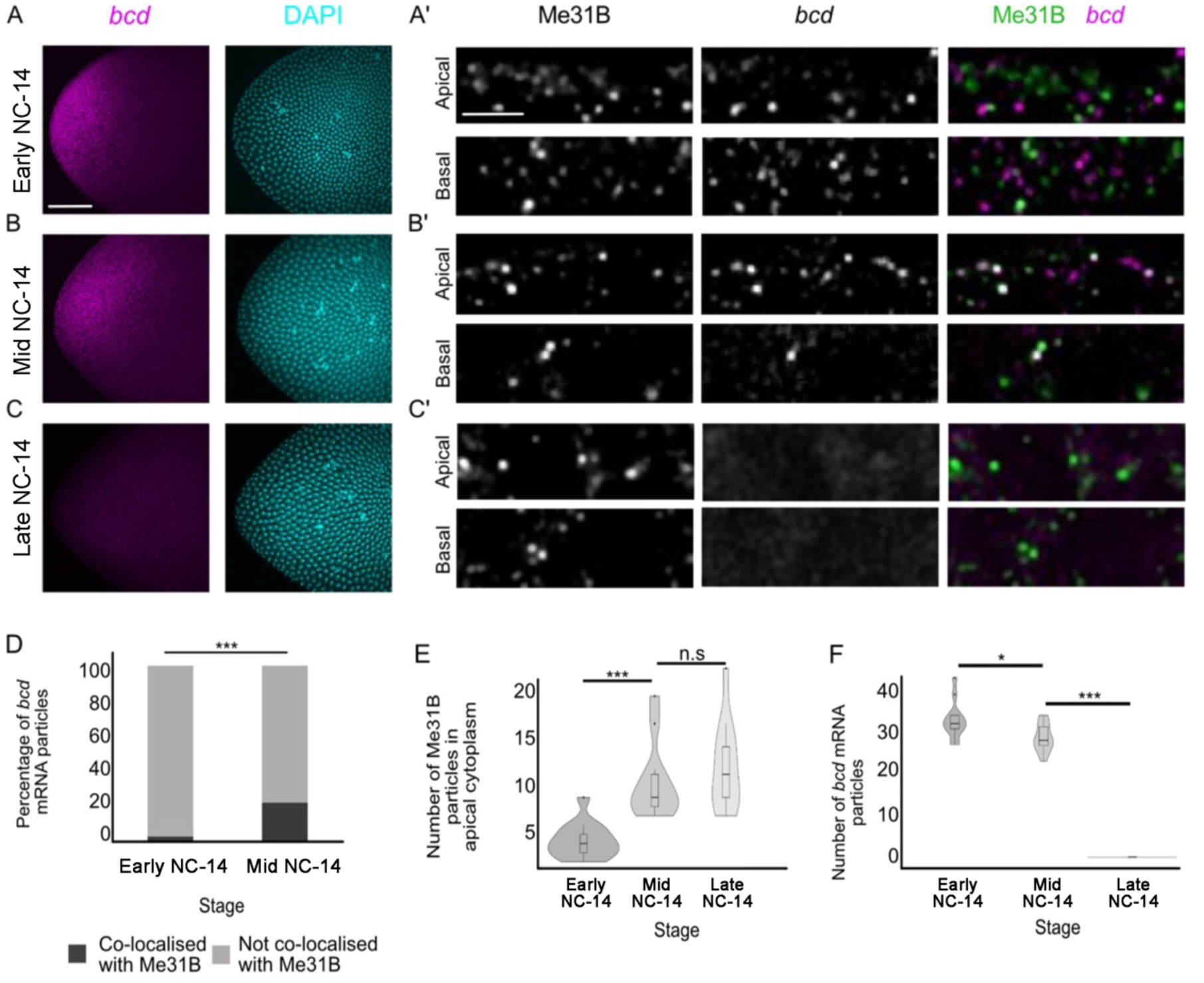
*bcd* re-associates with P bodies in n.c. 14, coinciding with their degradation. A-C: Representative fixed projected Z-stack images (from 20 embryos) showing *bcd* (magenta, smFISH) and DAPI (cyan) early n.c. 14, mid n.c. 14, and late n.c. 14. Scale bars: 50 µm. Aʹ-Cʹ: Representative fixed single plane images showing the apical side and basal side of early n.c. 14, mid n.c. 14, and late n.c. 14. Me31B::GFP (anti-GFP-nanobody), *bcd* (smFISH), and merge shown. N = 10 embryos. Scale bars: 10 µm; zoom 5 µm. D: Co-localisation between *bcd* and Me31B::GFP particles in the apical cytoplasm early and mid n.c. 14. N = 5 embryos. E: P body number in the apical cytoplasm between early n.c. 14 and mid n.c. 14. N = 5 embryos. F: Number of bcd particles present in the apical cytoplasm through early n.c. 14, mid n.c. 14, and late n.c. 14. N = 5 embryos. D-F: * *p* < 0.05; *** *p* < 10^*+^.

At mid n.c. 14, *bcd* was localised to the apical side of the nuclei (12). We found that it co-localised with embryonic P bodies, which were also enriched on the apical side (Figure 6Bʹ, E-F). As the embryo transitioned into gastrulation, we observed a significant loss of *bcd* while embryonic P bodies remained (Figure 6Cʹ-E-F). Together, this supports a model where, as the embryo transitions to zygotic control, maternal mRNAs in the embryo gather in embryonic condensates for degradation (79–81).

### Adapted SDD model of Bcd protein gradient formation

Formation of the Bcd morphogen gradient has been extensively modelled (11, 15, 82). Previous models typically considered the rate of Bcd production stayed constant during early embryogenesis. We then asked if considering the spatiotemporal pattern of *bcd* translation in the SDD model matched the observed Bcd gradient.

We considered the concentrations of *bcd* mRNA, shown as *[bcd]*, and the protein *[Bcd],* where the concentrations are functions of space and time, within a modified SDD model (Figure 7A):

**Figure 7:**
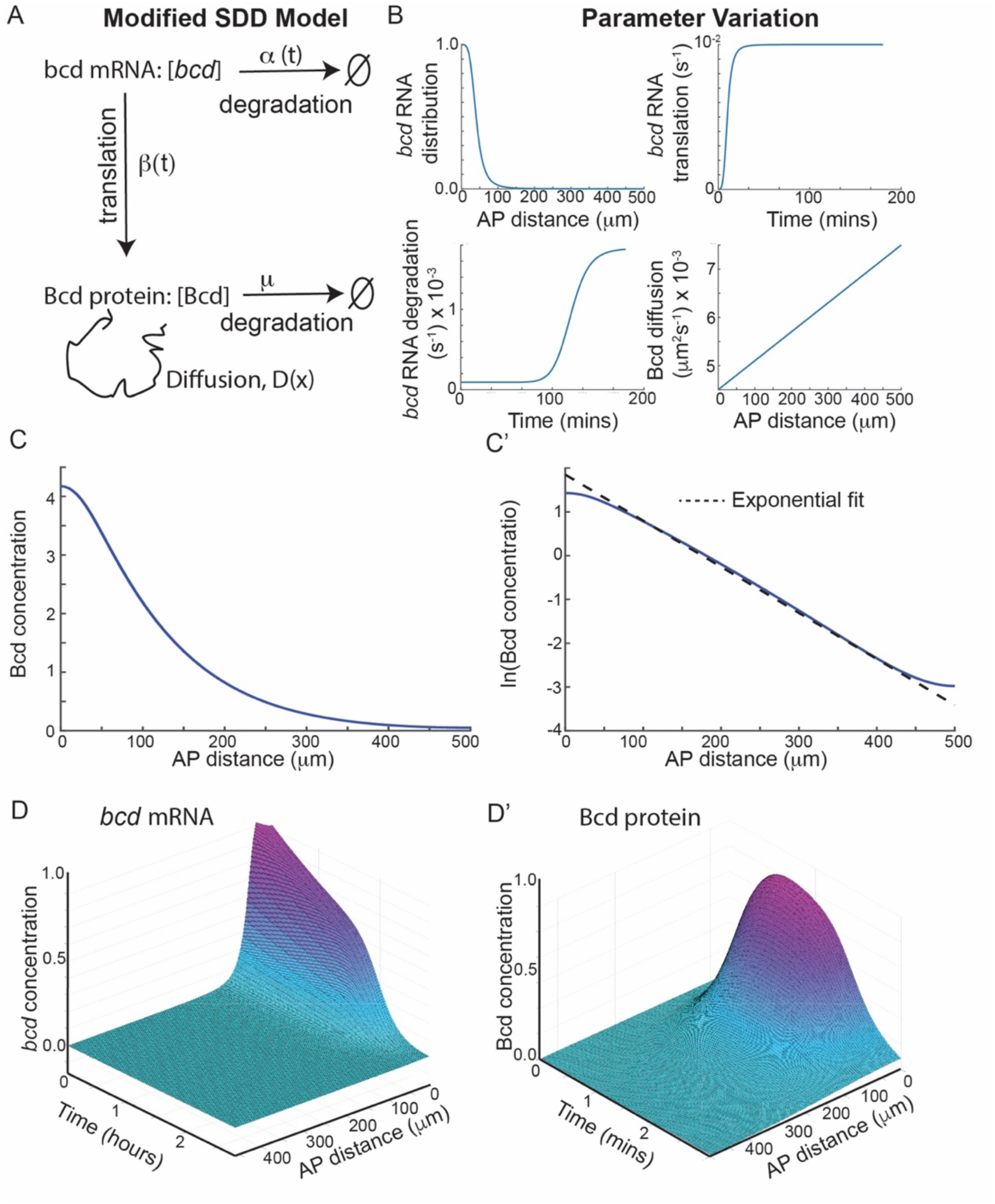
Modified SDD model of Bcd gradient formation. A: Model schematic, outlining parameters defining bcd and Bcd production, degradation and movement. B: Initial conditions (i) and parameter variation (ii-iv) over time or space within the model. C: Bcd concentration after 120 minutes in the simulation. C’: Same as C on log-plot, with an exponential fit shown with the dashed black line. Dashed line has decay length of 85 µm. D: Normalised concentration in space and time for *bcd* (D) and Bcd protein (Dʹ).

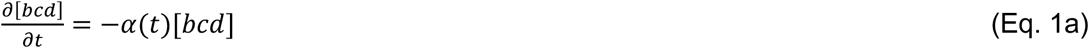

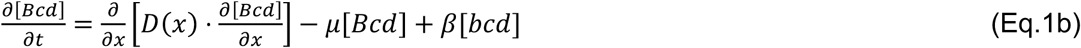

We defined time t = 0 to denote egg activation. The initial conditions for [*bcd]* and [Bcd] were given by:

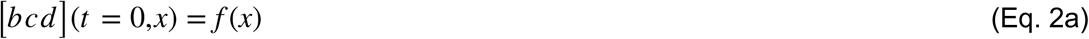

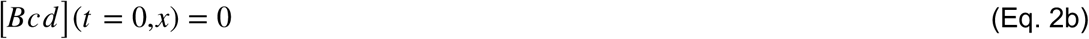

*bcd* mRNA was localised in the anterior pole with a form denoted by *f(x)*, (Eq. 2a), shown in Figure 7B(i). We incorporated a delay initially in translation to account for egg activation (Figure 7B(ii)). The rate of *bcd* degradation *α* (*t*) increased markedly in n.c. 14 (Figure 7B(iii)).

It was recently observed that the effective diffusivity of Bcd protein varies between the anterior and posterior pole (49). We included this through spatial variation in the diffusion coefficient *D(x)* (Figure 7B(iv)). We assumed that the rate of *bcd* translation after egg activation (*β*) and Bcd degradation (*μ*) remained constant. The degradation rate for Bcd has been calculated previously based on protein lifetime measurements (10).

Our modified one-dimensional SDD model generated an exponential-like gradient of Bcd in n.c. 13 (Figure 7C-C’). We emphasise that all the parameters in this model are constrained by experimental measurement, except for *β*, so it is not over parameterised (Table S5). We found a decay length of around 85-95 µm, consistent with experimental measurements (7, 10, 58). By including more realistic *bcd* mRNA translation, the overall behaviour of the Bcd protein gradient (Figure 7D-D’) is consistent with previous observations (58) and our quantification presented here.

## Discussion

We have provided a quantitative dissection of when and where *bcd* mRNA is translated in early *Drosophila* development. Translation is restricted until after egg activation by localisation within P bodies. Translation occurs in the anterior pole of the embryo and undergoes a marked decrease after n.c. 12. As embryogenesis continues, *bcd* becomes localised to the apical domain and is degraded, particularly at mid n.c. 14. This rapid degradation of *bcd* correlates with its localisation to P bodies during embryo cellularisation. These observations, along with recent quantifications of Bcd dynamics (49), support a modified SDD model of gradient formation.

In early embryos, Bcd protein is only observed in the anterior pole. This closely matches the *bcd* translation pattern. No significant translation was observed more than 50μm from the embryo anterior. These results argue that *bcd* translation does not occur in the oocyte and against a model of long-ranged protein gradient formation by a dispersed zone of production. Another model suggests that *bcd* is distributed through the embryo and hence can generate a long-ranged Bcd gradient without the need for diffusive transport (12, 83, 84). However, we observe most translation events occurred within 50 µm of the embryo anterior pole. Consistent with previous reports (58), we posit this is due to *bcd* localisation in the anterior region, rather than there being translational regulation of *bcd* in more posterior regions. This means that any graded form of *bcd* translation mostly happens in the anterior and that a long-ranged *bcd* translation gradient does not determine the Bcd protein gradient.

We noticed distinct differences in the apical-basal localisation of *bcd,* especially after nuclear migration to the embryo cortex. This change in apical-basal distribution correlates with a reduction in the observed *bcd* translation. However, it still needs to be tested if the localisation of *bcd* impacts its translation rate. The localisation of mRNAs within the microenvironment defined by each nuclear domain can impact translation in n.c. 14 (55), though whether a similar effect occurs earlier in development is unclear.

The rapid loss of *bcd* at n.c. 14 has been previously quantified (58) and here we discovered it also correlates tightly with localisation to P bodies (Figure 6). It remains unclear what determines the model of *bcd* degradation at n.c. 14, and if it is due to global degradation of maternal mRNA or a specific regulation of *bcd* mRNA (44). Testing for *bcd* degradation is challenging. One method is to infer degradation by examining spectral signal with different smFISH probes for the 5’ and 3’ UTRs (81, 85). However, the *bcd* 5’ UTR is very short, and we were unable to make effective probes. An alternative approach is to target the *bcd* instability element (BIE). This regulates degradation of *bcd* (78) and is located within the more accessible 3’ UTR.

It will be interesting to explore how differences in P bodies between the oocyte and embryo affects mRNA. P bodies appear to switch function, from shielding *bcd* from translation to enhancing *bcd* degradation. Prior to cellularisation, we found that Me31B condensates were insensitive to cycloheximide. However, after cellularisation begins, they become sensitised to cycloheximide (86), suggesting a change in their structure. A similar change in RNP function has been shown in *Drosophila* germ granules (87). Germ granules transition from protecting their mRNA contents to targeted degradation of specific mRNAs after germ cell specification. This is accomplished through recruitment of degradation factors as the granules grow through fusion. Decapping activity in these enlarged, functionally different germ granules lead the authors to call them P body-like. It is intriguing to consider that many RNP granules in *Drosophila*, and more broadly in *C. elegans, Xenopus*, zebrafish, will likely undergo changes in size, composition, and function throughout development. Relatedly, due to the conserved nature and wide-ranging downstream effects, it is intriguing to consider the possibility that Endos is a condensate regulator both within *Drosophila* eggs as well as more generally.

Over 40 years of work span from the original proposition of the SDD model for Bcd protein gradient formation. Here, we quantitatively assay the production of Bcd *in vivo*, which is arguably the last piece required to understand Bcd gradient formation. We see that the original SDD model, which assumed that dynamic parameters such as production rate and diffusion were constant in space and time, was too simple. However, with suitable modifications, like spatially varying diffusion and temporally altering translation, it can explain the formation of the Bcd gradient.

## Contributions

Project conception: ML, TES, TTW. Experimental design: TA, ELW, ML, TES, TTW. Experiments: TA and ELW with support from XS, PB, MV and ML. Analysis: TA, ELW, XS with support from AT, TES and TTW. Modelling: TES. Writing: TA, ML, XS, TES, ELW, TTW with all authors contributing to final version. Funding acquisition: ML, TES, TTW.

## Acknowledgements

We are very grateful to members of the Lagha, Saunders and Weil labs for support and comments on the project. We thank Jeremy Dufourt for advice on initial strategy for using Suntag with Bicoid and sharing plasmids (55). ML funding: ERC lightRNA2Prot, Marie Sklodowska-Curie Actions and CNRS. TES thanks core funding from the Mechanobiology Institute, Singapore and startup funding from Warwick Medical School. TTW thanks the Wellcome Trust Institutional Strategic Support Fund, University of Cambridge (grant number 097814) and Biotechnology and Biological Sciences Research Council Doctoral Training Partnerships studentship (ELW). ML acknowledges the Montpellier Resources Imaging facility (France-BioImaging) and the Drosophila facility (Biocampus). The lab of ML is funded by grants from the Fondation Bettencourt Schueller, the European Molecular Biology Organization Young Investigator program (EMBO YIP). The funders had no role in study design, data collection and analysis, decision to publish, or preparation of the manuscript.

## Methods

### Generation of SunTag *bcd* transgenic fly lines

*bcdSun32* was generated by replacing N-terminal eGFP of eGFP:Bcd-pCaSpeR4 construct (kindly gifted by Thomas Gregor (7)) by 32xSunTag repeats (kindly gifted by Mounia Lagha (55)). The 32xSunTag fragments are flanked by NheI and SpeI restriction sites at their N and C-terminal ends respectively by PCR, and then ligated into the eGFP:Bcd construct, replacing eGFP. Likewise, the first 10xSunTag repeats of 32xSunTag were ligated in the same way to generate *bcdSun10* construct (Figure S2A). Both Bcd SunTag constructs from the donor pCaSpeR4 construct were recombined into w+attB construct (addgene 30326) and flies were generated by phiC31 recombination into attP40 site of chromosome 2. The injection and generation of fly lines were done by BestGene Inc. USA.

The *bcdSun10* fly lines were crossed to the *bcd* null allele mutant, *bcd^E1^*. *bcdSun10*;*bcd^E1^* fly lines were viable, suggesting our construct can, at least partially, complement the function of endogenous Bcd. smFISH was carried out to test if *bcdSun10*;*bcd^E1^*effectively replaced the function of endogenous Bcd in the blastoderm. Stellaris smFISH probes with distinct fluorophores were used for SunTag and *bcd* coding regions of the *bcdSun10* mRNA. This enabled us to confirm that the only contribution of maternal *bcd* mRNA was from our *bcdSun10* in the *bcd^E1^* background (Figure S2). Co-localisation analysis in the early blastoderm revealed ∼90% of the mRNA were accounted for *bcdSun10* (Figure S2B). Further, the immunostaining of the blastoderm embryos with anti-GCN4 antibody revealed the translocation of the *bcdSun10* into the nuclei, unlike *bcdSun32*, eventually forming an anterior posterior gradient similar to endogenous Bcd (Figure S2C-E).

### Preparation of oocytes

Twenty female flies and ten male flies of the correct genotype were transferred into a vial containing fly food supplemented with yeast and were incubated at 25°C for forty-eight hours before dissection (88).

### Oocyte dissection

Fattened or aged female flies were anaesthetised, and the posterior tip of the abdomen was removed using a set of fine forceps. The ovaries were then removed from the open posterior end by squeezing the abdomen (88).

### Fly husbandry and preparation of embryos

Fly stocks were maintained at 21°C in vials that contained standard cornmeal starch agar fly food. To collect flies for experimentation or for genetic crosses, twenty females (virgin females were used for genetic crosses) and ten males were transferred into a plastic bottle containing yeasted cornmeal starch agar fly food at 25°C. Either virgin females or males from this generation were used for additional genetic crosses, or females younger than 10 days old were selected for experimentation. Collected embryos were dechorionated in 50% bleach for 2 minutes before excess bleach was removed with distilled water.

### Chemical and pharmacological treatments of oocytes

Hexanediol and sodium chloride: Dissected oocytes were incubated for 20 minutes in 5% 1,6-HD, 5% 2,5-HD, or 400 mM NaCl before immediate fixation. As controls, dissected oocytes were incubated in PBS or SIM for equal amounts of time before fixation.

*in vitro* activation of stage 14 oocytes (fixed): Dissected stage 14 oocytes were incubated in activation buffer (AB), for 30 minutes at room temperature. Activated oocytes were incubated in SIM for 0-30 minutes. To ensure that only activated oocytes were visualised, oocytes were treated with 50% bleach for two minutes. The activated oocytes were then fixed using the embryo fixation method. As a control, oocytes were incubated in SIM for the same time period. Drugs that inhibit the calcium wave: Dissected stage 14 oocytes were incubated in SIM that contained NS8593 for 30 minutes. The oocytes were then activated in vitro following the same steps as above, with all solutions containing the pre-incubation drug. As a control, oocytes were incubated in SIM (with no AB step) with the drug. Additionally, oocytes were incubated in SIM, followed by AB, followed by SIM with a 1:1000 dilution of either DMSO or ethanol.

### Fixation

Oocytes were incubated 1 ml of 4% paraformaldehyde for 15 minutes before immediate washing with 0.1% PBST (89). 10-20 pairs of ovaries were dissected and transferred to room-temperature Schneider’s Insect Medium (SIM). Ovaries were teased apart using dissecting probes until single ovarioles and oocytes were released.

Embryos were transferred into an Eppendorf tube that contains equal parts paraformaldehyde and heptane for 15-30 minutes, depending on the probes or antibodies used, before the paraformaldehyde phase was removed and replaced with 100% methanol. The sample was then shaken vigorously before all the solution was removed. Embryos were then rinsed with methanol and/or ethanol before being washed in 0.1% PBST (89).

### Single molecule fluorescence *in situ* hybridisation (smFISH)

#### Probe design

Custom 20 nucleotide Stellaris FISH Probes were designed against the 3ʹ UTR of *bcd,* coding region of *bcd*, and 32x SunTag by utilising the Stellaris RNA FISH Probe Designer and were the same as designed in (65) (Biosearch Technologies, Inc - www.biosearchtech.com/stellarisdesigner). 32x SunTag and *bcd* 3’UTR Probes were labelled with Quasar 570 and endogenous *bcd* is labelled with Quasar 670 were ordered for smFISH from Biosearch Technologies, Inc (Tables S2-4).

#### Buffers

Wash buffer was created by mixing the commercially available ‘Wash Buffer A’, de-ionised formamide and nuclease free water in a 2:1:7 ratio or was made using 2% sodium citrate buffer (SSC)+10% formamide. Hybridisation Buffer was created by mixing ‘Stellaris Hybridisation Buffer’ with de-ionised formamide in a 9:1 ratio.

#### Hybridisation

The hybridisation protocol was adapted from (55, 58). Briefly, fixed oocytes or embryos were permeabilised with 0.2% triton X-100 and washed in PBST. When necessary, embryos were blocked in PBST+RNase inhibitor+0.5% ultrapure BSA for 45 mins, before being washed in a ‘wash buffer’ solution. The samples were then pre-incubated in hybridisation buffer for two hours at 37°C. The hybridisation buffer was removed and replaced by the hybridisation buffer with smFISH probes for overnight incubation at 37°C, when using anti-GCN4 antibodies in conjunction with smFISH, primary antibodies were incubated overnight in the hybridisation buffer (90). The samples were then washed in hybridisation buffer before repeated washes in prewarmed wash buffer, when using anti-GCN4 antibodies in conjunction with smFISH, secondary antibodies were incubated with the final wash buffer wash, before being washed in 2X SSC. Finally, all samples were washed multiple times in 0.1% PBST.

#### Immunostaining

Nanobody labelling on oocytes or embryos, samples were blocked in a 4% BSA solution and then incubated with nanobodies, either GFP booster or RFP booster, in 0.1% PBST for one hour at room temperature. For anti-GCN4 immunostaining not in conjunction with smFISH, samples were incubated for two hours with the primary antibody, followed by washing with 0.1% PBST and incubation with the secondary antibody for two hours. After incubation, all samples were washed with 0.1% PBST.

Washed samples were transferred to a microscope slide and positioned using a dissecting probe. Excess PBST was removed before a single drop of Slowfade Diamond with DAPI or Prolong Diamond with antifade was added to the sample. A 22 x 22 mm coverslip was lowered onto the sample, and the coverslip was sealed with nail varnish and laid flat in the dark for 24 hours before imaging.

#### Imaging

All oocyte imaging was completed on an Olympus FV3000 inverted confocal microscope. Embryo imaging was performed on Olympus FV3000 or Olympus Spinning Disc systems. All images were taken using silicone oil 30x (0.95 NA) or 60x (1.4 NA) objectives. A region of interest was placed around either a whole single oocyte or embryo (30x objective) or a specific region of the oocyte or embryo (60x objective), Images were taken at a resolution of 1024 by 1024, or 2048 by 2048 pixels. 405, 488, 561 and 640 nm lasers were used. Laser power was adjusted to ensure that there was a full range of grey values and that the saturation of pixels was limited.

The images for the quantification of *bcd* foci in the embryo and their co-localisation to translating SunTag, were acquired on an Olympus FV3000 confocal microscope with the following conditions: 60x magnification, 4x zoom, 2 line averaging, slice thickness of 0.4 µm. The regions of high, moderate, and low dense *bcd* foci from the anterior domain and a very low *bcd* region in the posterior were imaged for a total thickness of 10-15 µm.

#### Fixed Imaging

Optimised step-size Z-stacks of fixed egg and embryos were collected; all Z-stacks were started at the cortex and extended 10-15 µm into the oocyte or embryo. For co-localisation studies, smFISH, and SunTag translation assays, where maximum resolution was imperative. Imaging parameters were optimised to improve resolution: 60 x objective used; pinhole: 1 airy unit; scan speed – 400 Hz; 2x line averaging; 2048 by 2048 pixels; and optimised Z-step (300 nm) (56).

### Image Analysis

Translation quantification: High-resolution image stacks were segmented to identify specific *bcd* foci. We used the trainable Weka segmentation plugin of ImageJ to train and classify the *bcd* foci. A classifier model created from a single image slice was applied to the Z-stack, thus classifying the *bcd* through the whole stack. The segmented particles were analysed using the Fiji particle analysis plugin that provided information on the size, shape, intensity and counts of all particles. The crowded particles that failed to segment properly, which was often the case within highly dense *bcd* regions, were not considered. We also excluded foci with low brightness. The segmented ROIs of *bcd* foci were applied to the anti-GCN4 channel. The mean intensities from anti-GCN4 channel were background corrected, and the positive mean values provided the number of co-localising SunTag to *bcd* particles *i.e.,* these particles were translating *bcd*. The plots were made in Graphpad Prism 8.0.

When automated methods were not accurate for detecting partial co-localisation (91), manual analyses were undertaken on images with only partial co-localisation. Images for both channels were thresholded, then an overlay of the threshold outline for channel one (*bcd*) was projected onto the channel two image (anti-GCN4). The number of particles in *bcd* channel that overlapped particles in anti-GCN4 channel was counted, and *vice versa*. To ensure that this partial co-localisation was not due to chance, images of one channel were inverted and the analysis was completed again.

### Statistical Analysis

n numbers: 10 to 20 females or 30 embryos (biological repeats) were used for each experiment and a global overview of the experimental results was acquired by visualising 20-30 oocytes or 30 embryos for each experimental condition unless stated otherwise. A smaller number of representative oocytes were subjected to high resolution imaging and analysis as described above. The “n” used in figures represents independent oocyte or embryo samples.

All statistical analyses were performed in R. For all statistical analyses, asterisks represent the following n.s. not significant * P<0.05, ** P<0.01, *** P<0.001.

Continuous data: Tests for normality were completed before proceeding with the appropriate parametric or non-parametric test. For comparison between two groups, an unpaired student t-test (STT) or a Wilcoxon signed-rank test (WSRT) was used.

Categoric data: A X^2^ test (CST) or Fisher’s exact test (FET) was used to determine if there were significant differences between groups. This depended on sample size Fisher’s exact test is preferred for small sample sizes (<50).

### Fly lines

Table S1 in the Supplementary Information outlines all lines used in this study.

### Reagents

Summary of reagents and tools used in this work

| Reagent | Supplier/ provider | Catalogue number | Concentration used |
| --- | --- | --- | --- |
| **Antibodies** |  |  |  |
| GFP Booster Alexa 488 | Chromotek | gb2AF488-100 | 1:1000 in 0.1% PBST |
| Rabbit GCN4 Antibody (C11L34) - Chimeric | Novus Biologicals | NBP2-81274SS | 1:200 in 0.1% PBST |
| Goat anti-Rabbit Cy3 | Thermofisher Scientific | A10520 | 1:200 in 0.1% PBST |
| Goat anti-Rabbit Alexa Flour 647 | Thermofisher Scientific | A21245 |  |
| **Buffers, chemicals, and pharmacological treatments** |  |  |  |
| Schneider's Insect Media (SIM) | Gibco | 21720024 | 1 X |
| Activation Buffer (AB) | Made in house: 3.3 mM NaH<sub>2</sub>PO<sub>4</sub>, 16.6 mM KH<sub>2</sub>PO<sub>4</sub>, 10mM NaCl, 50 mM KCl, 5% polyethylene glycol (PEG) 8000, 2mM CaCl<sub>2</sub>, brought to pH 6.4 with a 1:5 ratio of NaOH:KOH (260 mOsm) (Mahowald, Goralski and Caulton, 1983) | - | 1 X |
| Dulbecco's Phosphate Buffered Saline (without CaCl<sub>2</sub>) | Sigma Aldrich | D8537 | 1X |
| 1,6 Hexeandiol (1,6-HD) | Sigma Aldrich | 240117 | 5% in Schneider's Insect Medium or 1x PBS |
| 2,5 Hexeandiol (2,5-HD) | Sigma Aldrich | H11904 | 5% in Schneider's Insect Medium or 1x PBS |
| Sodium Chloride (NaCl) | Sigma Aldrich | S9888 | 400 mM in H<sub>2</sub>O |
| NS8593 | Sigma Aldrich | 492031 | Stock: 100 mM in ethanol<br>Final concentration: 100 μM in Schneiders Insect Medium or Activation Buffer (92) |
| Household Bleach | ESPO | 200-300-0011/1 | 50% in H<sub>2</sub>O |
| Paraformaldehyde | Thermofisher Scientific | J19943-K2 | 4% in PBS |
| Methanol | 32213 | Sigma Aldrich | 1 x |
| Heptane | Thermofisher Scientific | H/0106/PB17 | 1 x |
| Triton™ X-100 | Sigma Aldrich | X100 | 0.2% in PBS |
| Polyoxyethylenesorbitan Monolaurate (Tween-20) | Sigma Aldrich | P1379 | 0.1% in PBS |
| De-ionised Formamide | Sigma Aldrich | S4117 | 10%-30% in various FISH buffer solutions |
| 20 X SCC | Sigma Aldrich | S6639 | 2X-5X SCC in various FISH buffer solutions |
| Dextran sulphate | Sigma Aldrich | D6001 | 10% in various FISH buffer solution |
| 'Wash Buffer A' for Stellaris smFISH | LG Biosearch Technologies | Cat# SMF-WA1-60 | 20% in de-ionised Formamide and nuclease free H<sub>2</sub>O |
| 'Hybridisation Buffer' for Stellaris smFISH | LG Biosearch Technologies | Cat# SMF-HB1-10 | 90% with de-ionised Formamide |
| Hybridisation buffer for ImmunoFISH | Made in house (10% Formamide, 2xSSC, Ecoli tRNA (0.36mg/ml), VRC (2mM), 5% Dextran sulphate, ultra pure BSA(0.25mg/ml), stellaris probes, Primary antibody (1:500) |  |  |
| Tris EDTA (TE) Buffer | Made in house (10 mM Tris-HCl, 1 mM EDTA, pH 8.0) | - | 1 x |
| Slowfade Diamond Antifade Mountant with DAPI | Thermofisher Scientific | S36934 | 1 X |
| **Oligonucleotides** |  |  |  |
| (1) Custom Stellaris FISH Probes for the 3'UTR of *bcd* RNA labelled with a Quasar 570<br>(2) Bcd coding region Quasar 670<br>(3) 32xSunTag Quasar 570 | Stellaris | (See table 2.3 for list of smFISH probes) | Stock: 12.5 µM in TE buffer<br>Final Concentration: 50 nM in 0.25 ml Hybridization buffer |
| **Software and Algorithms** |  |  |  |
| Image J | Schneider et al., 2012 | https://imagej.nih.gov/ij/ |  |
| R Studio/ R Software | R Studio Team 2024 | http://www.rstudio.com/ |  |
| Graphpad Prism |  |  |  |
| **Other** |  |  |  |
| Standard Cornmeal Starch Agar FlyFood | Made in house by the Fly Facility, Department of Genetics, University of Cambridge | N/A | N/A |
| Olympus FV3000 Confocal Laser Scanning Microscope | Olympus | FV3000 | N/A |

