## Supplementary Information for "Dynamics of *bicoid* mRNA localisation and translation dictate morphogen gradient formation"

**Contents**

**Supplementary Figures**

**Supplementary Tables**

### Supplementary Tables

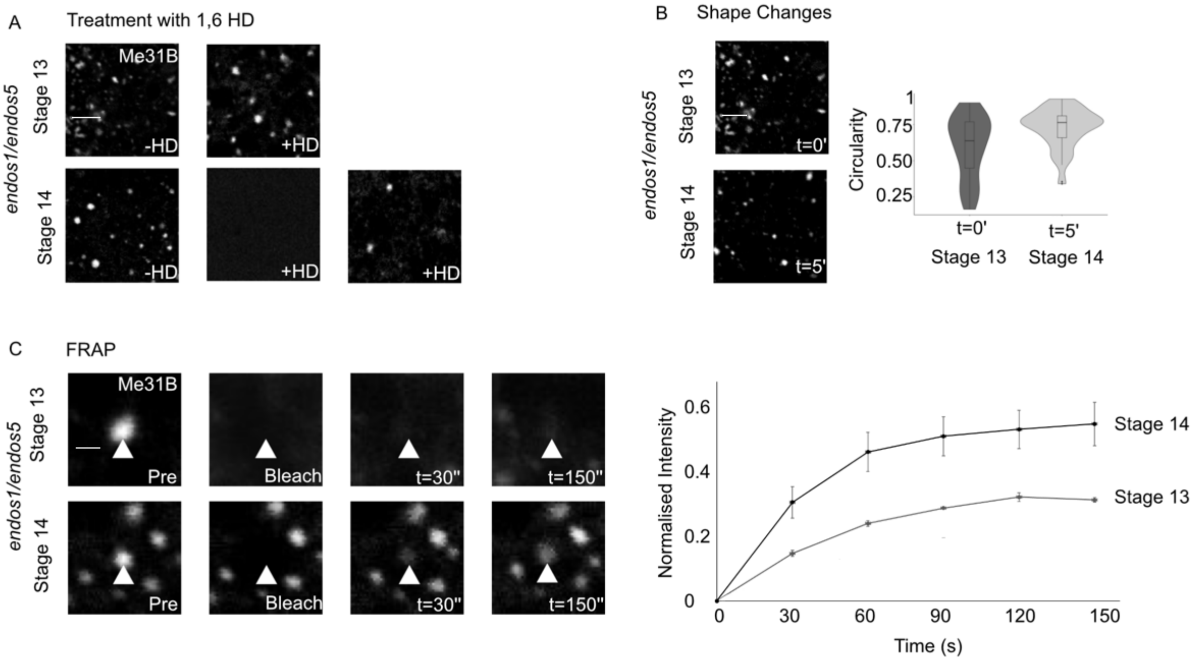


**Figure S1 (Related to Figure 2): P bodies in Endos mutant are less viscous after oocyte maturation**

A: Treatment of Endos mutant oocytes with 5% 1,6 hexanediol fails to disperse stage 13 oocyte P bodies within 20 minutes. The same treatment in stage 14 oocytes leads to almost complete P body dispersal consistent with a more liquid-like nature. N = 10 ooyctes for each stage.

B: Live imaging of Endos mutant oocytes during maturation followed by analysis of P body shape as measured by circularity. Wilcoxon signed-rank test, ** $p < 0.01$, N = 3 ooyctes for each stage.

C: Fluorescence Recovery After Photobleaching (FRAP) of P bodies in Endos backgrounds, shows minimal recovery in stage 13 oocytes. After oocyte maturation there is a higher recovery of fluorescence intensity in stage 14 oocytes, suggesting the condensates are less viscous as they exhibit more exchange with the cytoplasm. N = 5 ooyctes for each stage**.**

Scale bars: A-B = 5 μm; C = 2 μm

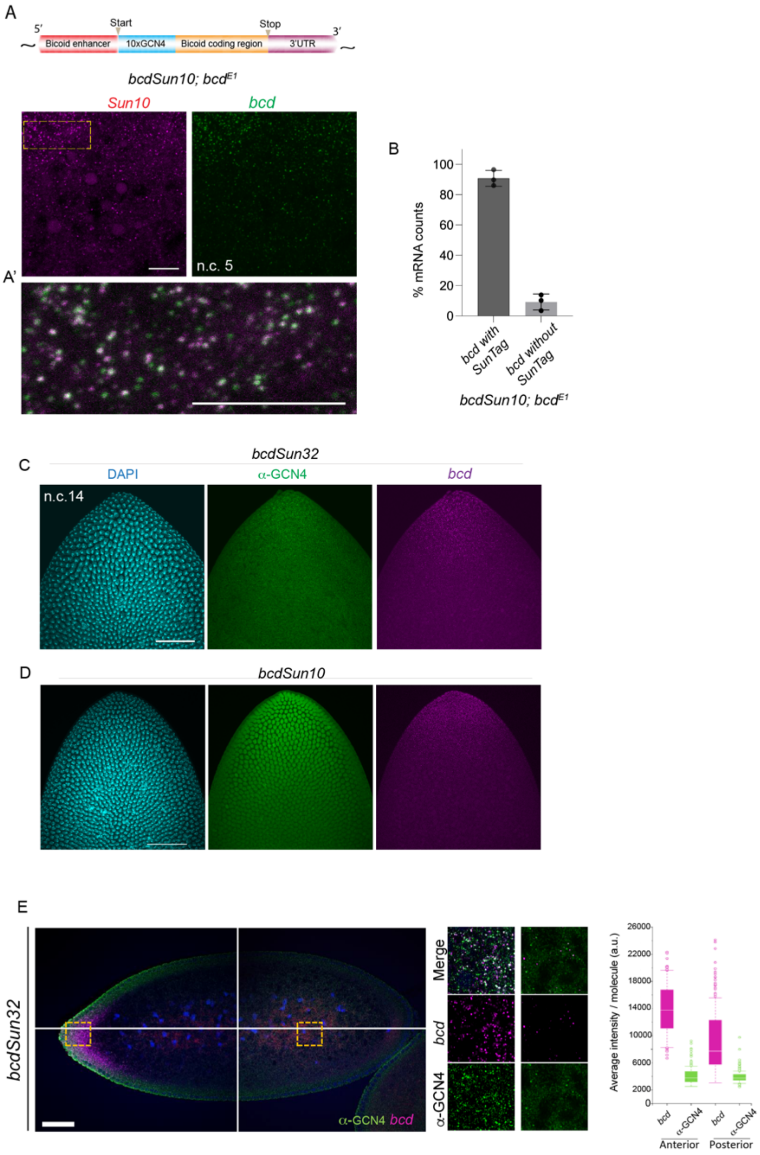


**Figure S2 (Related to Figure 3): *bcdSun10* rescues *bcd* null mutant**

A-A': (Top) Cartoon of *bcdSun10* construct. smFISH of SunTag (magenta) and *bcd* (green) regions of *bcdSun10* mRNA localised in the anterior domain of the early blastoderm of homozygous *bcd*$^{E1}$ mutant background. Yellow box magnified in A', showing the highly resolved *bcdSun10* mRNA particles.

B: Percentage of colocalisation of *bcd* and SunTag regions in the total *bcd* counts in the embryo's anterior pole (3 embryos). Error bars are SD.

C-D: Immuno FISH images of n.c.14 embryo. Embryos from *bcdSun32* (C) and *bcdSun10* (D) embryos are shown in upper and lower panels respectively (see Methods section for details).

E: Projected Z stacks of the anterior and posterior domains embryos expressing *bcdSun*32 transgene. *bcd* reporters and translation spots were detected by Immuno-FISH using Suntag probes and anti-GCN4 shown in magenta and green respectively. Quantification of the single molecule average intensity for bcd-reporters FISH and GCN4 IF signals in anterior and posterior domains (right panel).

Scale bars: A, A' = 10 µm; C-D = 50 µm

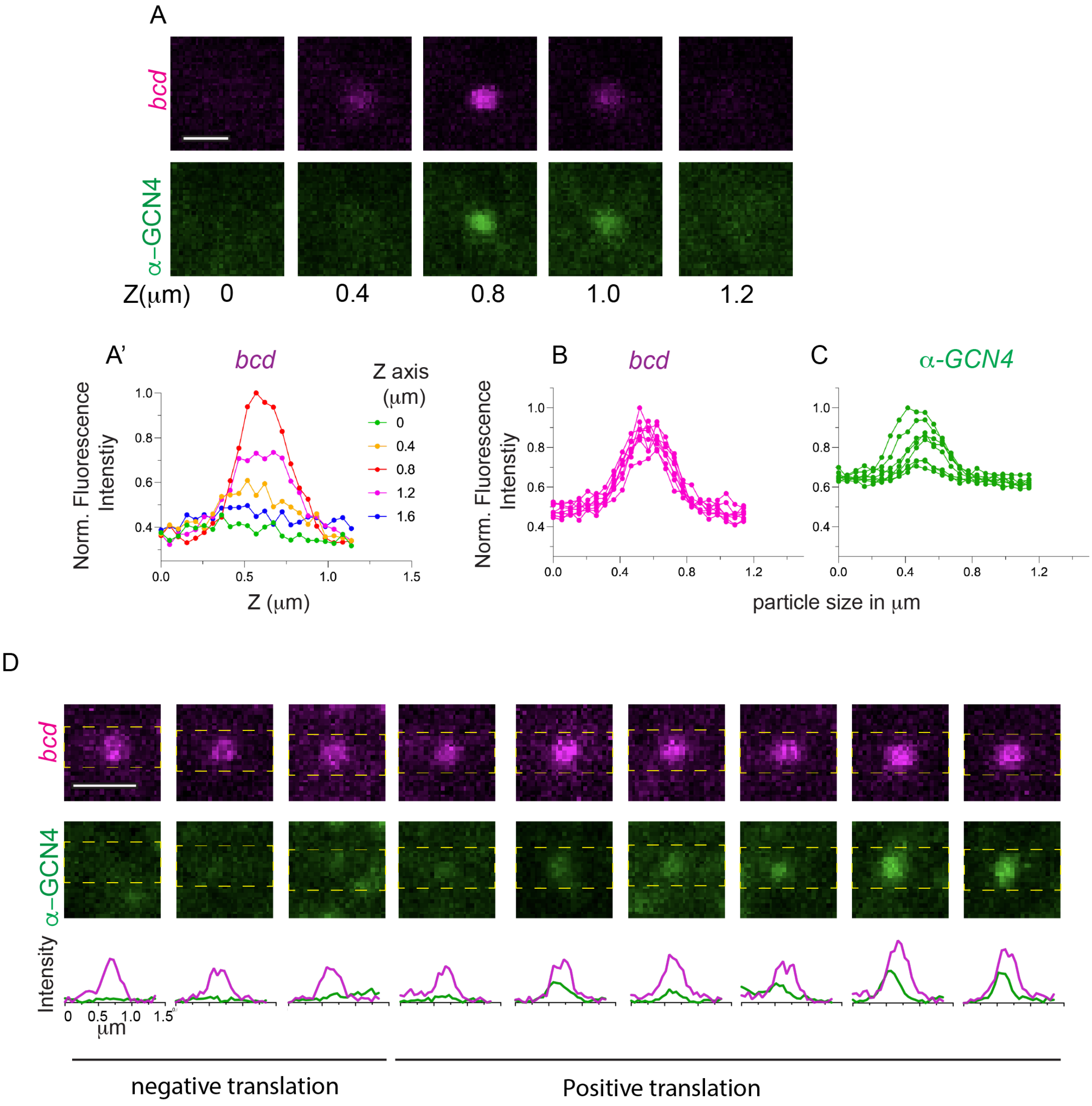


**Figure S3 (Related to Figure 3): Quantification of translating smFISH *bcd***

A: Z-slices of translating *bcd* (*bcd* (magenta), anti-GCN4 (Green)) from a *bcdSun10;bcd$^{E1}$* embryo at n.c.3. The *bcd* intensity from z-slices in A is captured in A'.

B, C: Maximum intensity of multiple *bcd* particles superimposed to show the overall spot size (Magenta, B) and variations in the intensity profile of SunTag (C, green). We find mean width of *bcd* spots is 0.7-0.8 μm. The intensity variations are shown in the bottom panel. Translating and non-translating *bcd* is classified based on the thresholded SunTag signal. Overall quantification is shown in B' and C'. Scale bar: A,D = 1 μm.

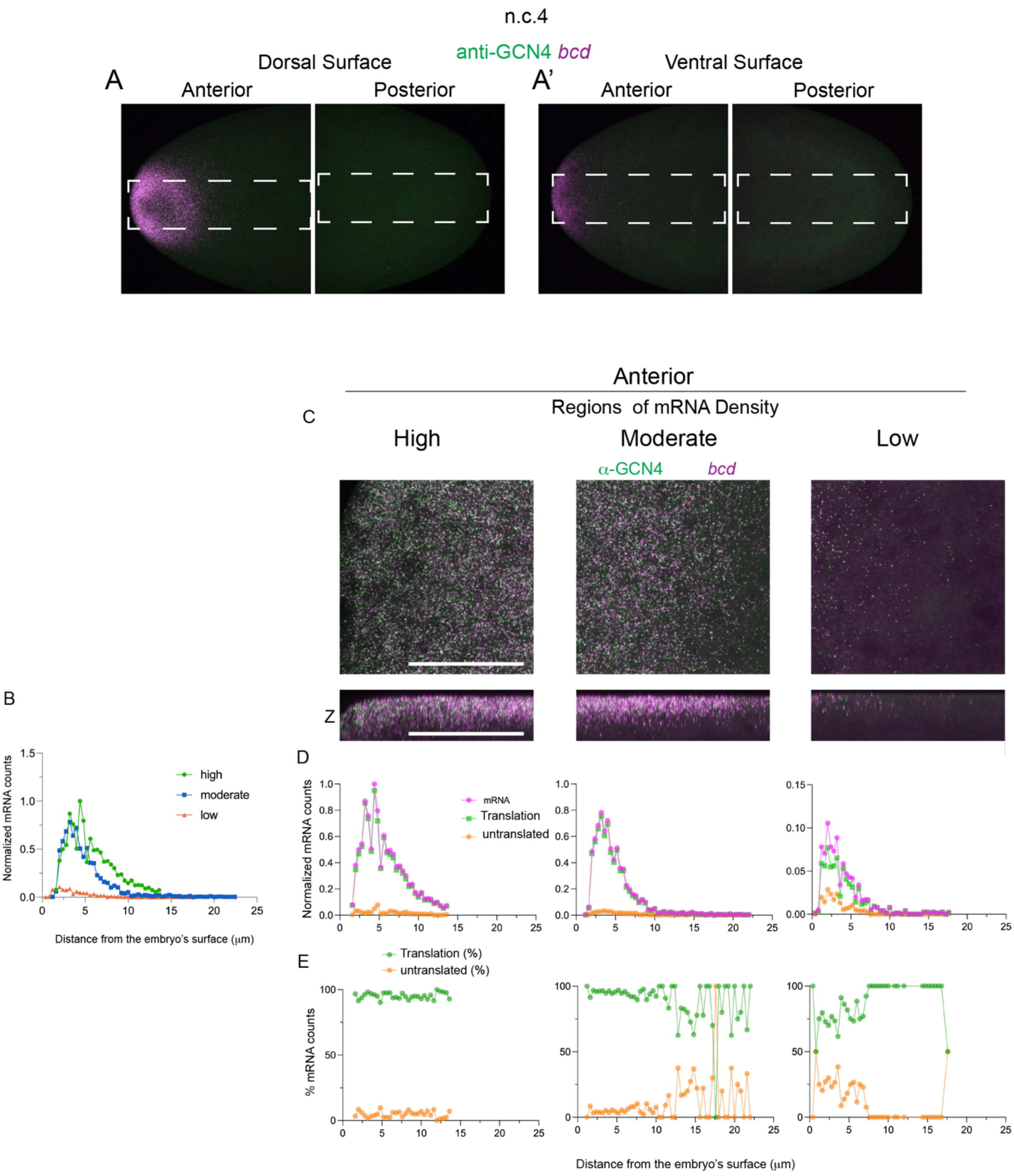


**Figure S4 (Related to Figure 3): Quantification of translation in the anterior domain of the embryo**

A: Immuno-FISH embryos at n.c.4 for *bcd* (magenta) and anti-GCN4 (Green). (B) The line plots of *bcd* spread on the Dorsal vs. Ventral surfaces.
B: Plot showing the distribution of *bcdSun10* counts from the surface-to-depth of the embryo at n.c.4 at the anterior domains where the mRNA are of high, moderate and low density. Counts are normalised to the total number of mRNA counts in the most anterior domain.
C: Maximum intensity projections of Z-slices and Z-cross sections at high, moderate and low dense *bcd* regions as quantified in B.
D-E: Normalised *bcdSun10* counts (D) and their percentages (E) undergoing translation at different positions from the embryo's cortical surface (2 embryos analysed). Counts are normalised to the total number of mRNA counts in the most anterior domain.
Scale bar: B = 50 µm.

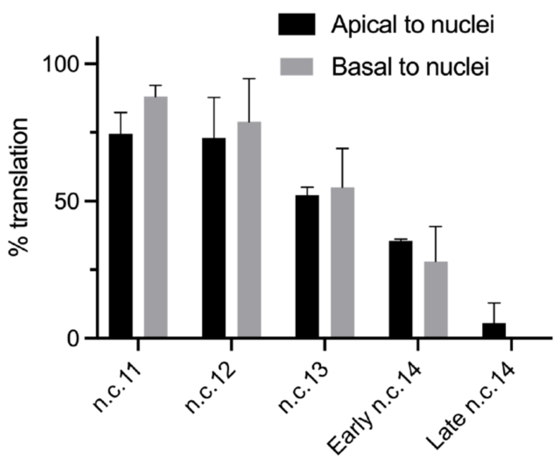


**Figure S5 (Related to Figure 4 and 5): quantification of translation of *bcd* apical and basal to the cortical nuclei**
Barplot showing variation in *bcd* translation localized apical and basal to the cortical nuclei. N = 3 embryos per nuclear cycle.

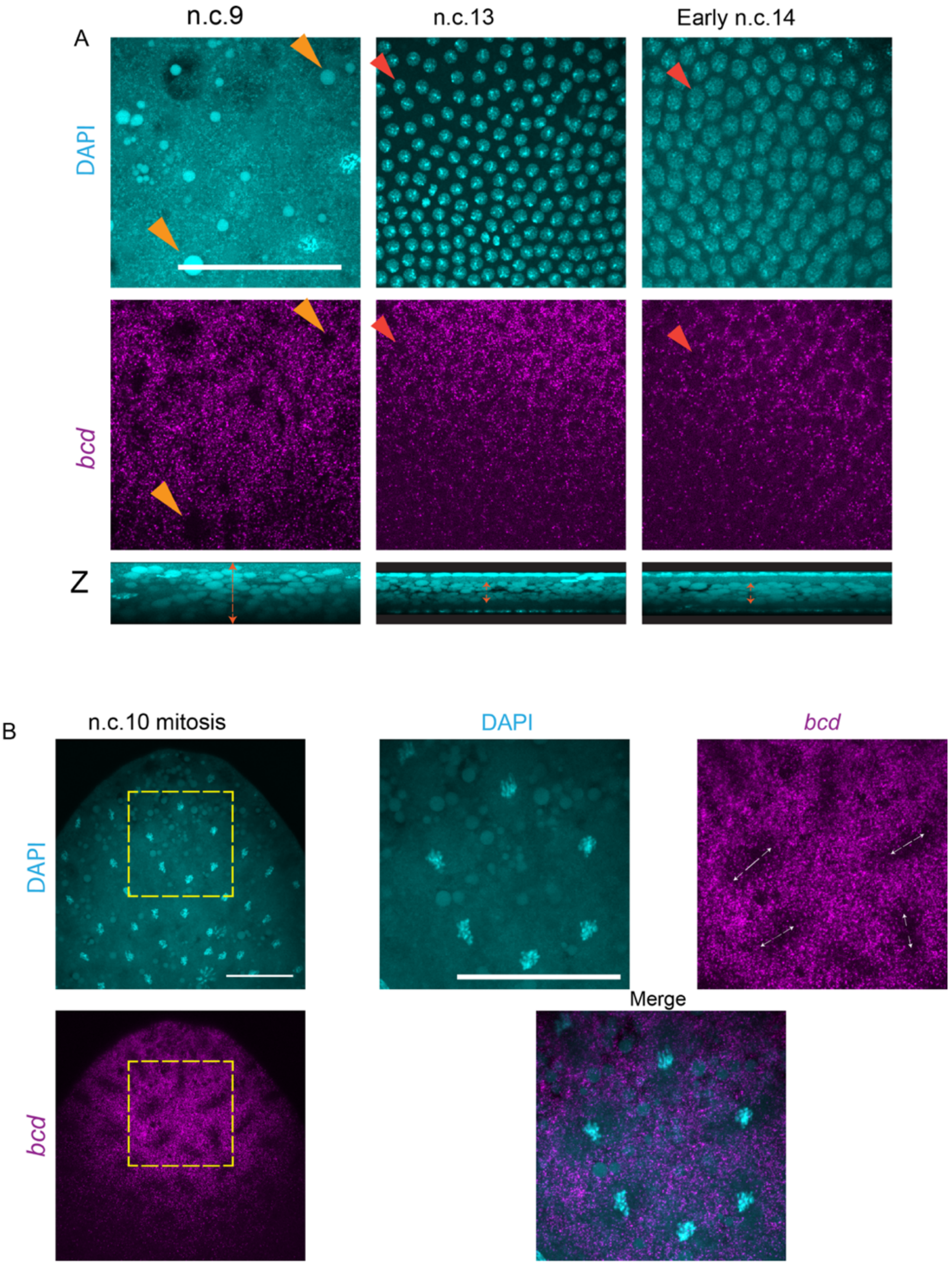


**Figure S6 (Related to Figure 4 and 5): Redistribution of *bcd* mRNA upon cortical migration and division of blastoderm nuclei**

A: Magnified Z-projected anterior regions of embryo cortex comparing stages n.c.9, n.c.13 and early n.c.14 stained for *bcd* (magenta) and DAPI (cyan). Orange arrowheads indicate lipid droplets, and the red arrowheads indicate nuclei. Double arrowheads indicate the embryo length along Z-axis where the lipids droplets are majorly spread.

B: n.c.10 mitotic embryo, with yellow boxes magnified on right. Double arrowheads indicate the direction of the nuclear division.

Scale bar: A-B = 50 µm

**Supplementary Tables**

| Fly line & genotype | Source and identifier |
|---|---|
| **Visualisation of proteins** | |
| **Me31B::GFP**<br>(Endogenous Tag) | (93)<br>BDSC: 51530 |
| **Suntag-Bcd**<br>W; 32xSuntag-Bcd/ Cyo<br>W;10xSunTag-Bcd;$bcd^{E1}$ | Generated here by Saunders lab |
| **Sc-FV-msGFP2**<br>nosEPr > Sc-FV-msGFP2 | Gift from Mounia Lagha<br>(91) |
| **Mutations and RNAis** | |
| **endos 1**<br>P(94); $endos^{1}$ /TM3, Sb | Gift from Hélène Rangone<br>(95) |
| **endos EY01103**<br>P(94); $endos^{EY01103}$ /TM3, Sb | Gift from Hélène Rangone<br>(95) |
| **endos EY01105**<br>P(94); $endos^{EY01105}$/TM3, Sb | Gift from Hélène Rangone<br>(95) |

**Table S1:** Summary of the *Drosophila* lines used in this work.

| smFISH probes |
|---|
| 5'-GAAACTCTCTAACACGCCTC-3' |
| 5'-ACAGTGGTTAACCTAAAGCT-3' |
| 5'-TGGTATTTGTACAATCAGGA-3' |
| 5'-CTTTCTACGCGTAGATATCT-3' |
| 5'-ACGGATCTTAGGACTAGACC-3' |
| 5'-AAACTTCCCTGGGAACCATT-3' |
| 5'-CTGCTGACTAGGCTAGTACA-3' |
| 5'-GATATGCACTGGAATCCGTG-3' |
| 5'-GAGTTAACTGGAGTATCACT-3' |
| 5'-AGCGTATTGCAGGGAAAGTA-3' |
| 5'-CACCCAGATACATCTAAGGC-3' |
| 5'-CATATTCCCGGGCTTTAGTG-3' |
| 5'-TGGCCTCAAATGTAACTGGT-3' |
| 5'-ACTTTCCATGGAATACGCTT-3' |
| 5'-ATTTCCGAAATGTGGGACGA-3' |
| 5'-AGAAGATTTTCTTGCTGGCT-3' |
| 5'-GTACAGTTTTTAGCTATGTC-3' |
| 5'-ATGAGATTACGCCCAAGAGA-3' |
| 5'-ATGTTCGATCTTTAAGGGTA-3' |
| 5'-ACACTTTGGCATAGCATAGA-3' |
| 5'-GCGCAAATGTTTGATTATGT-3' |
| 5'-TTGCTGACTATTCTTGGTCA-3' |
| 5'-ACAAATGGTCTGCATTGATT-3' |
| 5'-TGATAGTTATTCCGTTTGGC-3' |
| 5'-ATGCTCTTCTTAGTGATGTA-3' |
| 5'-ACTTGAGGCCTAACAGATTG-3' |
| 5'-ACAACATCAAAGGTGCAGCA-3' |
| 5'-ATTTACCCGAGTAGAGTAGT-3' |

**Table S2:** Custom Stellaris® FISH probes for the 3'UTR of *bcd* RNA (46).

| smFISH probes 32XSUNTAG |
|---|
| 5'- AGTCCTCTTCTTGAAGAGAG-3' |
| 5'- ATCTCTTACTTCATCGAGCT -3' |
| 5'- AGTCCACTTCTCGAGAATAG -3' |
| 5'- AGAACTTTTACTCCATCGCT -3' |
| 5'- AGACCACTTCTTAACGAGAG -3' |
| 5'- GATAGTGGAGCTCTTGCTTC -3' |
| 5'- CACCTCTTCTTGAAGACAGA -3' |
| 5'- TTAATGGTGGATCTCTTGCT -3' |
| 5'- CCACTTCTTAACGAGAGGTT -3' |
| 5'- GTAAACCTTTTACTTCACCG -3' |
| 5'- GTCGTTCTTGATAGTAGACC -3' |
| 5'- TTCATCGCTCTGACTTTTTT -3' |
| 5'- CGCTTCTTAACGAGAGGTTT -3' |
| 5'- ACCTCTTCTCAATGACTCAT -3' |
| 5'- AATCTTTTACTCCAGCGTTC -3' |
| 5'- TGAGGACAGATTCTTGATGG -3' |
| 5'- GTCTGAATTTTTTCCCTCAC-3' |
| 5'- AGGTTCTTGATGGTAGAGCT-3' |
| 5'- GATCCGACTTTTTTCCAAGA -3' |
| 5'- CGACGAGAGATTTTTGATGG -3' |
| 5'- CTTAATGGTAGACCTCTTGC -3' |
| 5'- CCAAGGCCACTTCTTGATAA -3' |
| 5'- AATGGTAGAGCTCTTACTCC -3' |
| 5'- CTCTTCTCAACGAGAGGTTT -3' |
| 5'- CTTCAACGCGCTGAATTCTT -3' |
| 5'- CTCTTCTTGACAACTCGTTT -3' |
| 5'- ATGGTAGAGCTTTTACTCCA -3' |
| 5'- TGGTAGAGCTCTTACTTCAC -3' |
| 5'- AGAGCTCTTACTTCAACGTG -3' |
| 5'- GTGAACCTTTTGCTTCAACG -3' |
| 5'- TCTTGATAGTGGATCTCTTG -3' |
| 5'- CCATCACCACTTCTTAATGA -3' |
| 5'- CTCTCCTCGATAACTCATTT -3' |
| 5'- TAAATCTCTTACTTCACCGG -3' |
| 5'- GGCCTCTTCTCAACAACAGT -3' |
| 5'- TGATTTCTTTCCAAGTCCTA -3' |
| 5'- CTTCTCGAGGAGAGTTTCTT -3' |
| 5'- CCGCTTCTTAACGAGAGATT -3' |
| 5'- CTAATTTTTTTCCATCACCC -3' |
| 5'- CTCCTTGATGAGAGGTTTTT -3' |
| 5'- TTCGCCACTCCTTAATGAAA -3' |
| 5'- AGTGGAGCTTTTGCTTCAAC -3' |
| 5'- GGCCACTTCTCGATAATAGA -3' |
| 5'- ATCTAATTTTTTCCCATCGC-3' |
| 5'- ACGATTCATTCTTGATGGTG -3' |
| 5'- GAATTTTTTCCCTAGACCTT -3' |
| 5'- TAGTGGAGCTCTTGCTTCAA -3' |
| 5'- TCTCTTACTTCATCGTTCTA -3' |

**Table S3:** Custom Stellaris® FISH probes for the 32xSunTag designed through stellaris probe designer version 4.2

| smFISH probes bcd coding region |
|---|
| 5'-GGCAGCGGATGATGGTAAAA-3' |
| 5'-AACTGAAGCTGCGGATGTTG-3' |
| 5'-AGGGATTTCGGAATTGTGGC-3' |
| 5'-CGTTCGCTCATCGAAAAGCA-3' |
| 5'-TGTAGTTGTAGTTTATCGCT-3' |
| 5'-CATCTGGTTGGGCAGATACG-3' |
| 5'-GAGGGAAAGACATCTGGCTT-3' |
| 5'-TAAAAGTGGTGCGGGTGCGA-3' |
| 5'-CAGCTCTGCTATTTGAGAGC-3' |
| 5'-TATCGTCCCTGCAGAAAGTG-3' |
| 5'-GCTAGTTTCGCTGACAGATC-3' |
| 5'-ATATCTTCACCTGGGCTGTG-3' |
| 5'-CGACGCCGACGGTTCTTAAA-3' |
| 5'-CTGATCCGATTGGATCTTGT-3' |
| 5'-TTTCATACCCGGCGAGAGAG-3' |
| 5'-AGCTAAGAGTCTGCAAGCTG-3' |
| 5'-GTGACGGAGTCAAAGCGTTG-3' |
| 5'-TGAATGACTCGCTGTAGTGC-3' |
| 5'-TCCATTGTAGTTGTAGTAGG-3' |
| 5'-CATGTGCATGTGACGATTGG-3' |
| 5'-ATTGACATTGGTCGACCCAG-3' |
| 5'-TTGCTTTTGCTGGAAGTCAA-3' |
| 5'-TTGAAGTTGTAGTCGGCCTC-3' |
| 5'-CGATCGCATGTAGTACGAGC-3' |
| 5'-GGTGTTAATGGCTCGTAGAC-3' |
| 5'-AGACTCGGACTTTCGTCATT-3' |
| 5'-AAGATCTGTAGCGTCGTCTT-3' |
| 5'-CTTGTCCAGACCCTTCAAAG-3' |
| 5'-TTCCTGCTAAGGCTCTTATT-3' |
| 5'-AAATGCCGCTCCACGATTTC-3' |
| 5'-GCGAAGGCTTGCCAAATTTG-3' |
| 5'-TGATATTGGTTCGATTCGCC-3' |
| 5'-TTGCATTATCGTATCCATCG-3' |
| 5'-CGTTCCGATGGGGATTATAC-3' |
| 5'-AGTAGGCAAACTGCGAGTTG-3' |

**Table S4:** Custom Stellaris® FISH probes for the bcd coding region designed through stellaris probe designer version 4.2.

| Parameter | Description | Value | Reference |
|---|---|---|---|
| $\alpha = \alpha_0 + \alpha_1 \cdot \frac{t^n}{t^n + t_0^n}$ | *bcd* degradation rate | $\alpha_0 = 1/180\ min^{-1}$<br>$\alpha_1 = 1/10\ min^{-1}$<br>$n = 12,\ t_0 = 120min$ | This work and (58) |
| $\beta \frac{t^m}{t^m + t_1^m}$ | *bcd* translation rate | $\beta = 10^{-2}s^{-1}$<br>$m = 4,\ t_1 = 10min$ | Fitted |
| $D(x) = D_0 + D_1 x/L$ | Bcd diffusivity | $D_0 = 4.5\mu m^2 s^{-1}$<br>$D_1 = 3\mu m^2 s^{-1}$ | (49, 97) |
| $L$ | Embryo length | $500\mu m$ | |
| $\mu$ | Bcd degradation rate | $1/35 min^{-1}$ | (10) |
| $f(x)$ | Initial *bcd* distribution | Figures 4 and S7 | This work |

**Table S5:** Model parameters used in Figure 7.
